# Thermal niche tracking in thirteen British temperate passerines

**DOI:** 10.64898/2026.04.24.720627

**Authors:** Ilaria Lonero, Mark J. Eddowes, Malcolm D. Burgess, James W. Pearce-Higgins, Albert B. Phillimore

## Abstract

Identifying how and why species vary in their ability to adjust to rapidly changing climates is a key challenge in ecology. While phenological shifts are well documented for birds and often studied in the context of tracking resource availability, less is known about the extent to which adjustments in phenology allow populations to track a consistent thermal niche. In particular, there has been little examination of how the extent of phenological thermal niche tracking compares over time versus space; a comparison that has the potential to inform on the underlying mechanisms. Here, we use data on breeding phenology derived from BTO Nest Record Scheme data, to examine the extent to which 13 passerine bird species track a consistent incubation thermal niche across years (both interannually and a year gradient) and along latitudinal and elevational gradients, and whether migrant and resident species differ in their tracking ability. Overall, we found support across species for partial tracking, with all species showing trends consistent with partial tracking across one or more axis, though for one species we could not reject the null hypothesis of no tracking. When we looked at average trends across species, we found significant tracking across interannual variation, latitude, and elevation, but not across a year trend. However, we found no evidence that tracking differs between residents and migrants, and for only a few species did we found evidence that species incubation thermal niche impacts on fitness. Taken together, our findings highlight the extent to which shifts in phenology can allow birds to track a thermal niche in a changing climate. The timing of a thermal niche provides a useful and widely-applicable yardstick to examine how changes in climate will impact on the abiotic conditions that populations experience.

## Introduction

One way climate change can impact on biodiversity is through a direct effect on the availability of favourable thermal conditions (Bellard et al., 2012; Pörtner & Farrell, 2008). A thermal niche defines the range of temperatures within which a species can survive, reproduce, and maintain physiological functions (Gvoždík, 2018; Magnuson et al., 1979). In addition to temperatures generally increasing with anthropogenic climate change throughout years, local temperatures vary substantially from year to year and across space. Seasonally, temperatures fluctuate considerably between summer and winter, with the amplitude of these fluctuations increasing with absolute latitude (IPCC, 2007). Spatially, temperatures decline by approximately 0.6–1 °C per degree of latitude from the tropics to the poles and by about 6.5 °C for every 1,000 m increase in elevation (IPCC, 2007). These temporal and spatial gradients provide axes along which species – especially those with narrower thermal limits – can track their thermal niche (Fredston et al., 2025).

One mechanism that may allow populations to track a broadly consistent thermal niche in the face of temperatures that vary across time and space is via phenological shifts. Phenological shifts involve changes in the timing of recurring biological events, such as leafing and flowering in trees, invertebrate emergence, or birds’ breeding and migration (Parmesan & Yohe, 2003; Roslin et al., 2021). For instance, in the spring, phenology considered at the population-level is advanced in warmer years (Charmantier et al., 2008; Cohen et al., 2018; Thackeray et al., 2016), and it is often later as one moves poleward or to higher elevation (Bell et al., 2019; Burgess et al., 2018; Hopkins, 1920). An impact of these shifts in timing is that they may contribute to ‘thermal niche tracking’, where populations maintain a relatively constant thermal niche by tracking spatial and temporal variation in its seasonal timing.

There has been considerable interest in how climate change may disrupt phenological interactions (Burgess et al., 2018; Charmantier et al., 2008; Vedder et al., 2013), with the shift in the timing of the resource often framed as a biotic ‘yardstick’ (Vedder et al., 2013; Visser & Both, 2005). In comparison, relatively few studies have explored the impacts on the abiotic environment an organism is exposed to, with the degree to which an organism tracks a consistent thermal niche representing an alternative abiotic yardstick. Laloë & Hays (2023) applied this concept in a global analysis of sea turtles, asking whether advancing nesting to cooler parts of the year could preserve their present-day thermal niche under a 1.5°C rise in sea surface temperature. Even under the strongest observed response—an 18-day advance per °C increase—nesting conditions were projected to warm by ∼0.6°C, indicating that phenological shifts alone cannot maintain thermal stability. Socolar et al. (2017), instead, reported that a 5–12 days advance in breeding phenology by Californian birds over the past century, reduced average nesting ambient temperatures by approximately 1°C, offsetting the increase in spring temperatures over this period. Similarly, López-Idiáquez et al. (2024) showed that whilst spring temperatures in Southern England have increased by 1.88°C over 59 years (1965–2023), Great tits (*Parus major*) maintained broadly stable temperatures within different reproductive phases.

Despite recent advances in understanding the role of phenological shifts in thermal niche tracking, substantial gaps remain. For instance, while temporally focused work has examined thermal niche tracking in relation to long-term temporal temperature gradients (Laloë & Hays, 2023; López-Idiáquez et al., 2024; Socolar et al., 2017), to our knowledge, previous work has not explored the ability of populations to track a thermal niche in the context of interannual temperature variation (Figure 1A). Whereas across space, most work on thermal niche tracking has focused on range shifts (e.g., Chen et al., 2011; Massimino et al., 2015; Neate-Clegg et al., 2024; Parmesan, 2006; Parmesan & Yohe, 2003), whilst the potential for thermal tracking to be achieved by phenological trends in space has been overlooked.

**Figure 1:**
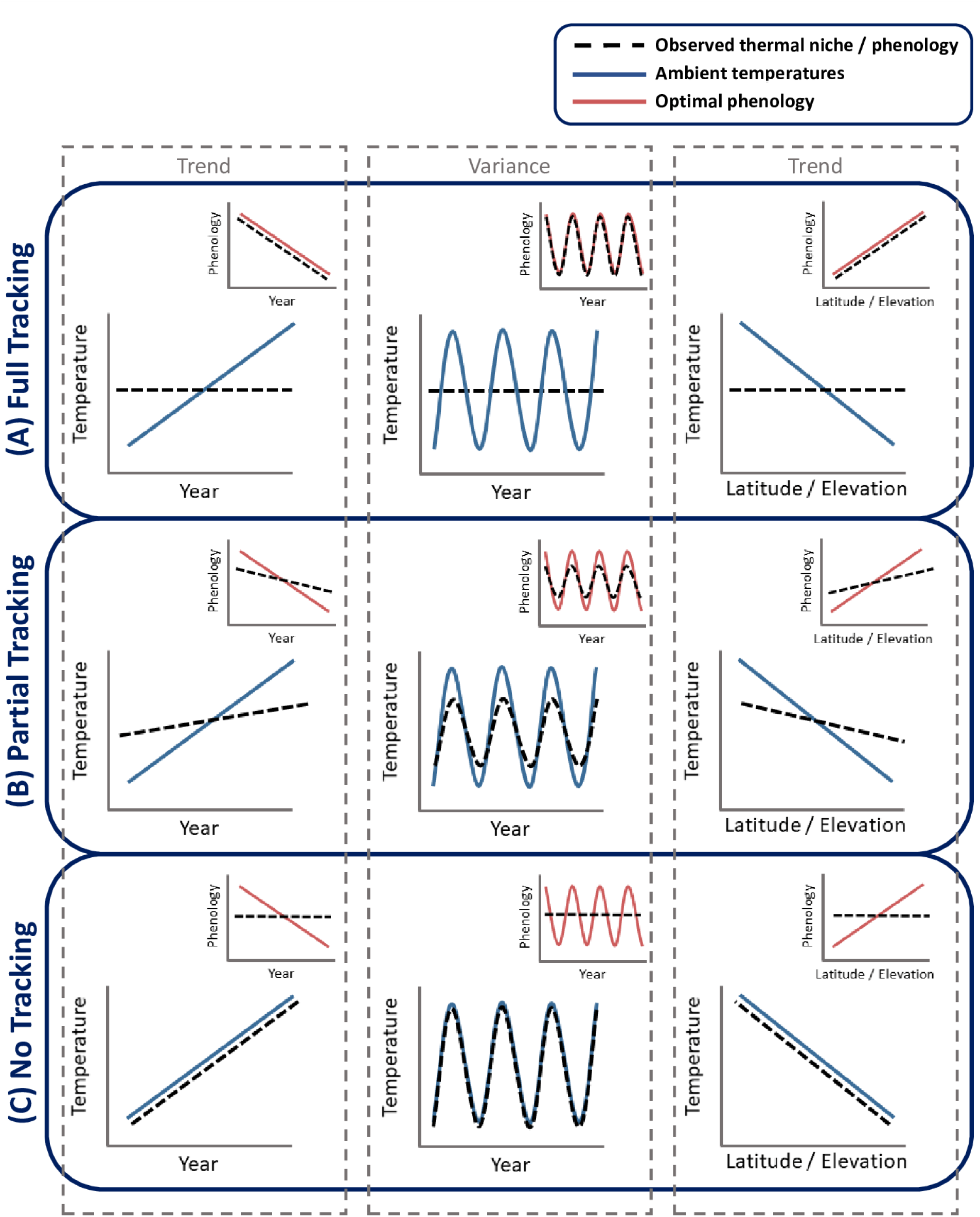
Thermal niche tracking scenarios. The schematic illustrates potential scenarios for thermal niche tracking across time (year) and space (latitude, elevation): (A) Full thermal niche tracking, (B) Partial thermal niche tracking, (C) No thermal niche tracking. Each scenario consists of two types of panels: temperature panels (large panels) and phenology panels (inset panels). The temperature panels display temporal (trend or interannual) and spatial (latitude or elevation trends) temperature. The blue line represents ambient temperatures, and the dashed black line shows temperatures relevant to the species’ thermal niche (the temperatures experienced by the species). The phenology inset panels illustrate the temporal and spatial shifts in phenology, with the red line representing the optimal phenology for full thermal niche maintenance, and the dashed black line representing realised phenology.

Understanding the extent to which a species tracks a thermal niche through phenology can provide insights into the impacts of rising temperatures on the timing and distribution of species. For instance, finding that a species exhibits strong thermal niche tracking across temporal and spatial gradients is consistent with phenological plasticity being the dominant contributor to timing variation (Figure 1A). Whereas, finding that a species exhibits thermal tracking across spatial gradients (Figure 1A, right panel) but not temporal gradients (Figure 1B,C, left and middle panels) can be explained if individuals are relocating or if there is local adaptation of timing (Phillimore et al., 2010) to maintain a thermal niche. Depending on the mechanism whereby tracking is achieved, leads to different predictions of how a population may respond to future climate change, specifically the contributions of plasticity, range shifts and genetic change.

Passerine birds have been a major focus in phenology studies due to the abundance of data on phenology of migration and breeding, evidence for thermal sensitivity of phenology (McLean et al., 2022; Saino et al., 2007) and their widely-documented phenological responses in relation to trophic interactions (Burgess et al., 2018; Vedder et al., 2013; Visser & Both, 2005). Across temperate passerines, timing of nesting coincides with the end of the winter and start of warmer spring conditions, though alongside this pattern, spring temperatures fluctuate considerably within and between years. A constraint for breeding birds is that ambient temperatures during the incubation period—the incubation thermal niche—can directly affect both parental energy demands and offspring fitness (Reid et al., 2000; Williams, 1996). Even slight variations in ambient temperatures during incubation (<1 °C) can affect embryonic development, particularly if prolonged periods fall below physiological zero (c. 26 °C), with potential consequences for hatchling success and fitness (DuRant, Hopkins, Carter, et al., 2013; DuRant, Hopkins, Hepp, et al., 2013; Hepp et al., 2006; Webb, 1987). For temperate breeding birds the timing of incubation (both in terms of onset and incubation window) is a trade-off between the costs and benefits of early breeding. While early breeding can be beneficial through reduced competition for nest sites (Dunn, 1977), lower predation risk (Nilsson, 1984), and improved reproductive success (Verhulst & Nilsson, 2008; Verhulst & Tinbergen, 1991), it also can incur higher energetic costs due to cooler average temperatures and greater risk of low temperature extremes (but also increased exposure to rainfall, e.g., Linden & Møller, 1989; Low & Pärt, 2009).

Prolonged exposure to cold ambient temperatures during incubation can lead to an extending of the incubation period, increasing the risk of nest predation (Martin et al., 2007) and energetic costs to incubating parents (DuRant, Hopkins, Hepp, et al., 2013). Given the temperature-dependence of invertebrate abundance (Macphie et al., 2023), cold temperatures also reduce food availability for foraging adults, with potential fitness consequences (Shipley et al., 2020; Winkler et al., 2013). Later incubation can also be costly, if eggs are exposed to elevated temperatures in warmer regions like deserts and the Mediterranean basin (McCowan & Griffith, 2021; Salaberria et al., 2014), or as a mismatch with the timing of food peaks, with potential consequences for chicks survival and breeding success (Visser et al., 2004; Visser & Both, 2005).

The ability of bird species to track an incubation thermal niche along different axes may depend on migratory behaviour. Compared with resident species, birds that migrate, especially long-distance migrants, face particular challenges adjusting their breeding phenology to track changing conditions in breeding areas, due to temperatures experienced at non-breeding sites tending to be uncorrelated with interannual variation in temperatures on the breeding grounds (Both & Visser, 2001; Hulme, 2001; but see Selonen et al., 2021 for some evidence of weather links between stop-overs and arrival sites), limiting the ability of migrants to plastically adjust their arrival time (Finch et al., 2014; but see Ockendon et al., 2013 for significant advancement in migrants’ egg laying dates). A lack of advancement in arrival phenology shortens the period between arrival at breeding sites and egg-laying, placing a constraint on the ability of migratory birds to plastically advance their breeding time *in situ* (Both et al., 2006; Both & Visser, 2001; Strode, 2003). These constraints likely explain why long-distance migrants show weaker temperature sensitivity in their arrival and nesting phenology compared with short-distance migrants and residents (Samplonius et al., 2018; Usui et al., 2017). However, this does not imply that migrants are less affected by temperature per se: indeed, there is evidence that their breeding success can be more sensitive to spring warming than in residents (Pearce-Higgins et al., 2015), though the extent of such mismatch effects remains debated (Franks et al., 2018). Intriguingly, evidence is coming to light that advances in breeding timings of some migratory species may have a genetic component (Helm et al., 2019; Lamers et al., 2023; Lonero et al., 2024). Across space, species that are less able to respond plastically to spring conditions may rely more on alternative mechanisms to track the thermal niche via natal (or breeding) dispersal to suitable climates (Lamers et al., 2023) or via local adaptation (Phillimore et al., 2010).

Migratory species may have greater propensity than residents to track a thermal niche through natal dispersal to territories that present suitable thermal conditions in a particular year (Bonnet-Lebrun et al., 2021; Gómez et al., 2016; Shutt et al., 2022). Migratory birds also typically have later phenology than residents, so generally experience warmer incubation temperatures, thereby reducing risks associated with cold conditions in cooler climate, and exacerbating risks from high temperatures in warmer climates. Consequently, selection pressures related to thermal niche tracking may be weaker for migrants in cool temperate climates compared to residents, potentially making temporal and spatial tracking less critical.

Our aim is to examine the ability of passerine birds breeding in the UK to track a thermal niche over time and over space. We extend the approach of López-Idiáquez et al. (2024) to study thermal niche tracking along a temporal gradient by considering interannual tracking, and tracking in space across latitudinal and elevation gradients. We apply our approach to 13 forest passerine species and first assess the ability of each species to track an incubation thermal niche over different gradients, and then estimate the average level of tracking for each gradient across species. We follow this with a test of the hypothesis that migratory species are less able to track the thermal niche across years and interannual variation compared with residents, but are better at tracking their thermal niche across latitude and elevation. Finally, to test the hypothesis that there is selection on the incubation thermal niche, we examine whether low (and high) incubation temperatures are associated with lower fitness.

## Materials and methods

### Data

#### Nest monitoring data

We used British Trust for Ornithology (BTO) / Joint Nature Conservation Committee (JNCC) Nest Record Scheme (NRS) breeding records for 13 passerine species that are largely single-brooded in the UK, which breed primarily in woodland habitats and that provide at least 1,000 derived phenology records. Focusing on single-brooded woodland species ensured consistency in comparing thermal niche tracking, while also targeting a group particularly vulnerable to climate change impacts on breeding timing and success. The NRS is a long-term citizen science project that systematically collects data on bird breeding attempts across the UK, relying on volunteers to monitor nests through multiple visits (Crick et al., 2003). Although the scheme spans from 1939 to the present, our study used data from 1980–1990 to 2023 to align with the availability of corresponding temperature records (see below the paragraph on *Temperature data),* with fewer records for the earliest years because of imprecise nest location data. Our study focused on five resident species: Blue Tit (*Cyanistes caeruleus*), Chaffinch (*Fringilla coelebs*), Great Tit (*Parus major*), Long-tailed Tit (*Aegithalos caudatus*), and Nuthatch (*Sitta europaea*); and eight migratory species: Garden Warbler (*Sylvia borin*), Pied Flycatcher (*Ficedula hypoleuca*), Common Redstart (*Phoenicurus phoenicurus*), Sedge Warbler (*Acrocephalus schoenobaenus*), Spotted Flycatcher (*Muscicapa striata*), Whinchat (*Saxicola rubetra*), Willow Warbler (*Phylloscopus trochilus*), and Wood Warbler (*Phylloscopus sibilatrix*). The sample size for each species dataset is showed in Table 1.

**Table 1:** Sample sizes for each species dataset used in the *Selection on the thermal niche* (left) and *Thermal niche tracking* (right) analyses. For each species, the number of records is shown for every data-editing step (S0–S13), with final sample sizes highlighted in bold. Editing steps: S0 = combined NRS and Met Office dataset including all nest records with corresponding temperature data; S1 = removed records with invalid or implausible first egg or hatching dates (e.g. due to inconsistent years of nest visit); S2 = retained records with first laying date uncertainty ranges ≤ 20 days and removed missing values; S3 = removed statistical outliers; S4 = deleted error measurements; S5 = excluded records with missing clutch size measurement; S6 = calculated incubation start and end dates, removing missing values; S7 = derived minimum incubation temperatures and laying date deviations; S8 = defined fixed (null) windows and measured their minimum temperatures; S9 = corrected a coordinate error for Pied Flycatcher; S10 = extracted elevation data from AWS DEM; S11 = filtered invalid OS grid references; S12 = removed nest records located outside UK land polygons; S13 = retained records within 0–1,000 m elevation, with complete year, latitude, and elevation values, and mean-centred variables.

|  | Sample size prior to selection analyses |  |  |  |  |  |  |  | Sample size prior to thermal tracking analyses |  |  |  |  |  |
| --- | --- | --- | --- | --- | --- | --- | --- | --- | --- | --- | --- | --- | --- | --- |
| Species | S0 | S1 | S2 | S3 | S4 | S5 | S6 | S7 | S8 | S9 | S10 | S11 | S12 | S13 |
| Blue Tit | 127,020 | 126,527 | 94,518 | 93,705 | 84,684 | 84,684 | 84,684 | **84,405** | 84,405 | 84,405 | 84,405 | 84,251 | 83,716 | **83,680** |
| Chaffinch | 8,412 | 8,409 | 6,403 | 6,351 | 6,279 | 6,279 | 6,279 | **6,273** | 6,237 | 6,237 | 6,237 | 6,016 | 5,889 | **5,889** |
| Great Tit | 87,141 | 86,887 | 54,525 | 53,852 | 49,851 | 49,851 | 49,851 | **49,696** | 49,696 | 49,696 | 49,696 | 49,541 | 49,185 | **49,177** |
| Long-Tailed Tit | 4,537 | 4,479 | 3,058 | 3,017 | 2,817 | 2,817 | 2,817 | **2,804** | 2,804 | 2,804 | 2,804 | 2,795 | 2,765 | **2,765** |
| Nuthatch | 6,159 | 6,120 | 4,774 | 4,725 | 4,241 | 4,241 | 4,241 | **4,237** | 4,237 | 4,237 | 4,237 | 4,237 | 4,222 | **4,222** |
| Garden Warbler | 1,462 | 1,462 | 1,424 | 1,412 | 1,363 | 1,363 | 1,363 | **1,361** | 1,361 | 1,361 | 1,361 | 1,361 | 1,361 | **1,361** |
| Common Redstart | 5,850 | 5,845 | 5,222 | 5,154 | 4,951 | 4,951 | 4,951 | **4,941** | 4,941 | 4,941 | 4,941 | 4,941 | 4,938 | **4,938** |
| **Pied Flycatcher** | 38,204 | 38,182 | 36,601 | 35,810 | 34,076 | 34,076 | 34,076 | **3,4072** | 34,072 | 34,071 | 34,071 | 34,071 | 34,060 | **34,060** |
| **Sedge Warbler** | 1,888 | 1,888 | 1,781 | 1,779 | 1,744 | 1,744 | 1,744 | **1,692** | 1,692 | 1,692 | 1,692 | 1,658 | 1,488 | **1,488** |
| **Spotted Flycatcher** | 5,671 | 5,667 | 4,994 | 4,985 | 4,884 | 4,884 | 4,884 | **4,875** | 4,875 | 4,875 | 4,875 | 4,800 | 4,784 | **4,761** |
| **Whinchat** | 2,569 | 2,567 | 2,443 | 2,429 | 2,369 | 2,369 | 2,369 | **2,369** | 2,369 | 2,369 | 2,369 | 2,364 | 2,357 | **2,357** |
| **Willow Warbler** | 6,462 | 6,462 | 5,847 | 5,757 | 5,673 | 5,673 | 5,673 | **5,642** | 5,642 | 5,642 | 5,642 | 5,605 | 5,516 | **5,516** |
| **Wood Warbler** | 2,406 | 2,404 | 2,348 | 2,324 | 2,281 | 2,281 | 2,281 | **2,281** | 2,281 | 2,281 | 2,281 | 2,281 | 2,277 | **2,277** |

The NRS records provide information on nest location and inspection dates where counts of eggs and/or live chicks are made. Recorders typically visit active nests every 5–7 days. The NRS then generates estimates of minimum and maximum laying dates (i.e., first egg dates) through back-calculation, using egg counts, nestling developmental stages, laying rates, incubation periods, and nestling periods (Crick et al., 2003). Clutch size is estimated as the total number of eggs at the end of the laying period (Crick et al., 2003).

#### Nest monitoring data preparation

All data processing was conducted in R (v.4.2.2, R Core Team, 2025). Our focus was on the incubation thermal niche. Each species’ dataset was subset to include only nests where warm eggs or hatching were observed, indicating nests that had incubated eggs. We estimated the incubation start date for each nest as the sum of the midpoint for laying date (midpoint between minimum and maximum estimated values) and clutch size (i.e. we do not include egg-laying pauses or pauses prior to incubation, as we lack information on these), thus assuming that birds lay one egg per day and start incubation the day the last egg is laid. The incubation end date was determined as the incubation start date plus the species-specific average incubation period (BTO, 2025), allowing consistent comparisons across species. We used species-specific average incubation periods to avoid complexities arising from incomplete or imprecise observational data. We restrict our focus to observations where the difference between minimum and maximum laying date estimates was ≤20 days. We filtered the NRS dataset to include only records without calculation errors detected from BTO validation programs. Nest locations were reported at either 100m or 1km accuracy. For each species, we identified and removed invalid references using the R package *sgo* (version 0.9.2, Lozano Ruiz, 2022). Eastings, northings, latitude, and longitude were extracted from valid references through package *sgo*, but a further ∼3,000 coordinates located off land were excluded based on the *rnaturalearth* (version 1.0.1, Massicotte & South, 2023) coastline dataset. For each valid square, we derived latitude and longitude coordinates as well as 50-km grid cell identity from easting and northings, and elevation (through package *elevatr*; version 0.99.0, Hollister, 2023). Hatching probability was measured using a binary metric, with nests containing at least one alive chick on day 5 post-hatching classified as successful (nest success = 1) and nests with no alive chicks considered unsuccessful (nest success = 0). We used this binary classification rather than counting the number of hatched eggs to reduce the impact of observer error when counting broods. Survival of nests to day 5 was used as an approximation for hatching probability to focus on impacts on fitness incurred during the incubation phase, with the aim of minimising the effects on fitness arising during later nesting stages.

#### Temperature data

Daily minimum temperature data were spatially linked to individual nest records for the months in which the estimated laying date and incubation period occurred—generally March–June for resident species and April–August for migratory species. These temperature data were interpolated at 1 km resolution across the UK, using temperature from Met Office stations (Perry et al., 2009; www.metoffice.gov.uk/climatechange/science/monitoring/ukcp09/). This approach provided a detailed temporal and spatial framework to evaluate incubation conditions. Minimum temperatures during incubation are a key determinant of incubation success in temperate passerines (DuRant et al., 2010, 2014), and this underpinned our decision to focus on the minimum of the daily minimum temperatures during incubation.

### Statistical Analysis

Our analysis consisted of three steps. First, we examined how well each species tracks its incubation thermal niche in both time (year gradient and interannual variation) and space (latitude and elevation gradients). Second, we compared the average thermal niche tracking between resident and migratory species. Finally, we quantified selection on the incubation thermal niche, by analysing the relationship between the minimum temperature experienced during incubation and our hatching probability approximation.

#### Thermal niche tracking

To assess how each species tracks its incubation thermal niche, we built on the framework described by López-Idiáquez et al. (2024) quantifying the thermal change experienced by a population during a particular phenological stage, and comparing this to a null expectation of the thermal change that would be experienced if the phenology was invariant. For analysis of the observed thermal gradients and fluctuations in time and space, we fitted Bayesian Generalized Linear Mixed Models (GLMMs) using the MCMCglmm package in R (version 2.34, Hadfield, 2010). Minimum temperature during incubation for each nest record was used as the response variable. We included year, latitude and elevation as fixed effects. All the fixed effect variables were mean-centred prior to analysis. For two species—Garden Warbler and Sedge Warbler—the range of nest elevations was limited (see Supplementary Material), potentially constraining our ability to detect meaningful elevation-based thermal niche tracking. We therefore caution against overinterpreting elevation-related results for these species and excluded them from both our estimates of average tracking across species and resident–migrant comparisons. Random effects were year, 50 km grid cells, the interaction between year and 50 km grid cells, and 5 km grid cells, to account for temporal, spatial, and spatiotemporal pseudoreplication. Our focal model parameters were the gradients of (i) year, (ii) latitude and (iii) elevation, plus (iv) the year random effect. The year random term was included to capture short-term annual variability in incubation temperature (interannual year changes in temperature), distinct from the long-term trends estimated via the year fixed effect.

To generate a null expectation for trends/variances in environmental temperature for each species, we considered minimum temperature during fixed windows of the same length as the species-specific average incubation period. The number of fixed windows was set to span the range in incubation periods observed in the nest records across all individuals of the same species (Figure S1), with more windows for species with a bigger range in incubation period (e.g., 7 windows for Blue Tit, and 11 windows for Chaffinch). For each fixed window, we extracted the 1km and year specific temperatures for every nest record for the species-specific incubation duration and calculated the nest specific minimum temperature. The fixed window model included the minimum temperature during the window as response variable, and the same fixed and random terms as outlined above. After estimating parameters for each fixed window, we then concatenated the posteriors for the different fixed windows to generate a null that accounts for both uncertainty in the parameter estimates and heterogeneity in temperature trends/variances across different stages of the spring.

We used default priors in MCMCglmm for the fixed effects (normal distribution with mean = 0, and a large variance), an inverse Wishart distribution (V = 1, nu = 0.02) for the residual, and parameter expanded priors for the random effects. Models were run for 400,000 iterations with a burn-in of 20,000, and a thinning interval of 100. We checked the models by visualisation of trace files for convergence (see Supplementary material). For all models, we required a minimum effective sample size of 1000 for each parameter.

#### Comparing observe and fixed window temperature trends

Thermal niche tracking would be evidenced where incubation temperatures remained stable over time (year trend and interannual variance) and space (latitudes and elevations), even when the average fixed window temperatures—calculated across the posterior of the different fixed windows—showed temporal or spatial variation. The average fixed window slopes (or variances) served as the null expectation for the temperature change expected if there was no phenological shift and thermal niche tracking. An observed term (slope or variance) = 0 would be consistent with perfect thermal niche tracking, an observed term (slope or variance) = the null term, would be consistent with no niche tracking, whereas an observed term > 0 but < the null term would be consistent with partial tracking. The above comparisons were all made for each species across the observed and null posteriors. For each species, we also quantified a tracking metric using the model posterior, as the ratio between the absolute observed trends in incubation temperatures and the absolute trends in the fixed window temperatures. A tracking metric of 0 indicates perfect thermal niche tracking, while a value of 1 indicates no tracking (note that a value of 1 could arise if the observed gradient in temperature was of the same magnitude but opposite direction to the fixed window gradient). Values in the range 0-1 would correspond to partial tracking and values >1 would arise if the temporal or spatial change in temperature exceeds what would be expected if incubation timing was held constant (Figure S2).

#### Cross-species trends in tracking performance

To examine whether tracking is generally stronger over certain axes we estimated the median tracking metric across all 13 species, and repeated this across a posterior of estimates to obtain a median and 95% CIs. The pairwise difference between the posteriors estimated for the four axes provides a test of whether tracking differs between them.

#### Thermal niche tracking comparisons between resident and migratory species

After calculating the tracking metric for each effect of interest (year-gradient, interannual, latitude, elevation) for each species, we estimated the median of the tracking metric across the posterior for residents and across migrants separately. We then used the posterior of the cross-species medians to infer a median and 95% CIs for residents and migrants. As a metric of comparison, we finally calculated the difference between the resident and migrant posterior metrics, and we extracted the median difference (and 95% CIs). If the credible intervals of the difference did not overlap zero this would indicate a significant difference in tracking between residents and migrants.

We conducted comparisons both including and excluding Garden Warbler, Common Redstart, Spotted Flycatcher, and Willow Warbler as we observed patterns in the lay date versus reproductive success relationship that appear to be consistent with a substantial proportion of second broods. In a single brooded species, we expected hatching success to either show a unimodal peak or to decline with later laying date. However, for these migratory species we observed a bimodal peak in hatching success, with peaks in early and late nests, which we interpreted as consistent with having a second brood.

#### Selection on the thermal niche

To test for directional or stabilising selection on the incubation thermal niche, we evaluated the effect of minimum incubation temperatures (as both a linear and a quadratic term) on our approximation of hatching probability for each species. We used Bayesian Generalized Linear Mixed Models (GLMMs) fitted through the MCMCglmm package (Hadfield, 2010), with hatching probability treated as a binary response. We examined the evidence for two hypotheses regarding the effect of the incubation thermal niche on avian fitness: (1) stabilising selection where fitness is highest at intermediate temperatures and (2) directional selection where fitness is lowest at cold temperatures and increases with incubation temperature. Laying date is typically negatively correlated with breeding success in birds (Radchuk et al., 2019), so to control for this we included annual laying date deviations (calculated within 1km grid cells) as linear and quadratic fixed effect terms. Models also included a fixed effect of year, which we scaled for the analyses on Blue Tit. To account for spatial, temporal, and spatiotemporal pseudoreplication, we included random effects for 50 km grid cells, year, their interaction, and 5 km grid cells. We also included a random slope-and-intercept term that allows the relationship between laying date deviations and hatching probability to vary across years and 50 km grid cells.

MCMCglmm priors, iterations and burnin were as described above.

R version 4.2.2 was used for all data preparation and analyses (R Core Team, 2025).

## Results

Based on the fixed windows (null) trends, the 13 UK passerine species experienced a non-significant increase in ambient spring (spanning windows between 72-242 ordinal days, or 13^th^ of March – 30^th^ of August) temperatures of 0.03 °C/year, equating to ∼1.5 °C over 50 years (95% CI: –0.01 to 0.06; Table S1), with substantial interannual variability (2.2 °C, 95% CI: 1.02 to 3.99; Table S2). Temperatures decline by 0.33 °C/°N across latitude (95% CI: – 0.44 to –0.20; Table S3) and by 0.3 °C/100m elevation (95% CI: –0.4 to –0.3; Table S4).

Based on these spatial gradients, the 1.5°C temporal increase would correspond to a poleward shift of approximately 4.5°N (95% CI: 3.4–7.5), translating to ∼500 km (in the range of ∼375–830 km), and an upward elevational shift of ∼500 m (in the range of ∼375–750 m) to fully track climate change.

### Thermal niche tracking across the year thermal gradient

Across years, the only species that experienced a significant directional change in incubation temperature were Spotted Flycatcher (0.03 °C/year, 95% CI: 0.01 to 0.04, Table S1) and Whinchat (0.02 °C/year, 95% CI: 0.001 to 0.04, Table S1). The average fixed window temperature was non-significant for the remaining species, ranging from -0.01 to 0.02 °C/year (Table S1). We found no significant differences between the observed year slopes and the corresponding average fixed window slopes (Table S1, Figure 2A). All species showed partial thermal tracking of the year gradient, with tracking metric medians ranging between 0 and 1 (Table S5, Figure 2B), but the credible intervals were wide in all cases, arising from considerable variation in the across-year temperature gradient experienced during different periods in spring. The species that exhibited the strongest tracking, in order, were Chaffinch, Nuthatch, Long-tailed Tit, Pied Flycatcher, Common Redstart, and Blue Tit (Table S5, Figure 2B).

**Figure 2:**
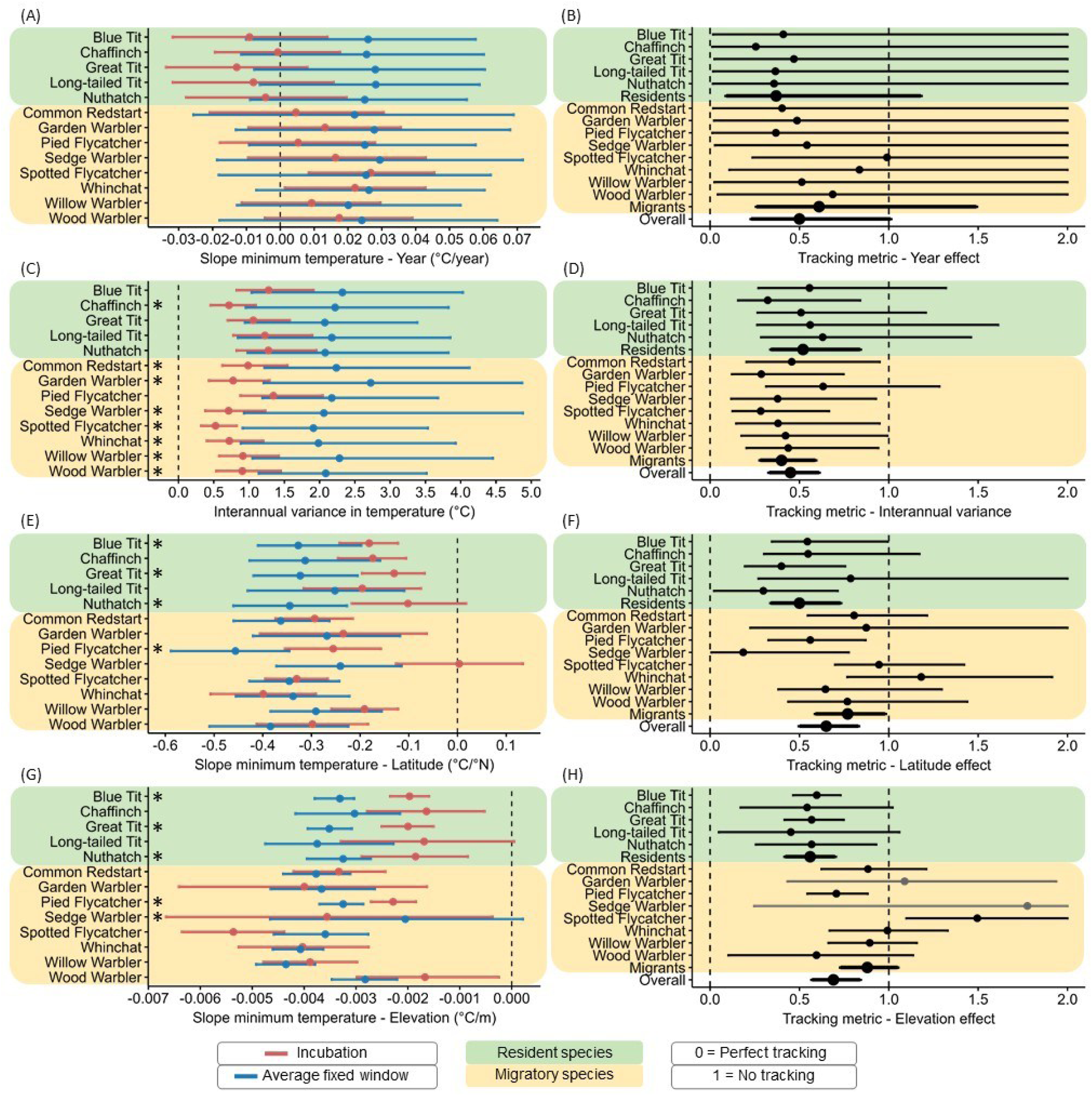
Trends in incubation temperatures (red; left column) and fixed window temperatures (blue; left column) and thermal niche tracking metrics (0 = perfect thermal niche tracking, 1 = no thermal niche tracking; right column) for thirteen passerine species in the UK, analysed across (A, B) years, (C, D) interannual variations, (E, F) latitude, and (G, H) elevation. The green panels illustrate trends for resident species, while the yellow panels illustrate trends for migratory species. The slopes, along with their posterior medians and credible intervals (CI) are shown. In the left plots (A, C, E, G), the slopes, along with their posterior means and credible intervals, are presented for both incubation and fixed window effects. Incubation slopes indicate changes in temperatures experienced during the incubation period. Fixed window slopes represent temperature changes during fixed time intervals. An incubation slope that overlaps zero is consistent with maintaining a consistent incubation temperature across the specified gradient. Asterisks next to the species names highlight significant differences between the incubation slopes and fixed window slopes. In the right plots (B, D, F, H), the thermal niche tracking metrics are presented as the ratio between the observed trends in the species’ incubation temperature and the trend in fixed window temperature. A tracking metric (slope and CI) of 0 indicates perfect thermal niche tracking, while a value of 1 indicates no tracking. Metrics (slope and 95% CIs) between 0 and 1 are consistent with partial thermal niche tracking. The overall tracking metric reported in each plot represents the tracking metric estimated across all species for each gradient, while the residents and migrants tracking metrics are measured across each species group. The tracking metric across elevation is in grey for Garden Warbler and Sedge Warbler since they were excluded from the comparative analyses due to their limited ranges of elevations (see Table S7), potentially constraining our ability to detect meaningful elevation-based thermal niche tracking. The x-axis was truncated in panels B, F, and H to better highlight values closer to the 0–1 range; however, the confidence intervals for some species extend beyond the displayed range (see Table S5).

### Thermal niche tracking across the interannual thermal gradient

Incubation temperatures varied from year-to-year for all species, as did the average fixed window temperatures, with the variance estimates consistently lower for the observed incubation temperatures (Table S2, Figure 2C). Significant differences between observed and null variance were found for 7 of the 13 species: Chaffinch, Garden Warbler, Common Redstart, Sedge Warbler, Spotted Flycatcher, Whinchat, Willow Warbler, and Wood Warbler (Table S2, Figure 2C).

All species showed partial interannual tracking, with tracking metric medians ranging from 0.3 to 0.6 (Table S5, Figure 2D). However, only for Chaffinch, Garden Warbler, Common Redstart, Sedge Warbler, Spotted Flycatcher, Whinchat, and Willow Warbler was the upper CI < 1 (Table S5, Figure 2D).

### Thermal niche tracking across the latitude thermal gradient

Incubation temperatures did not change significantly across latitude for Nuthatch and Sedge Warbler (Table S3, Figure 2E), indicating a degree of consistency in the incubation thermal niche, whereas for other species incubation temperatures decreased significantly with latitude. Average fixed window temperatures showed significant decreases with latitude for all species (Table S3, Figure 2E), with the slopes generally steeper than for the incubation temperatures (Table S3). Significant differences between observed incubation trend slopes and average fixed window slopes were detected for Blue Tit, Great Tit, Nuthatch, and Pied Flycatcher (Table S3, Figure 2E).

Most species showed partial tracking across latitudes, with tracking metric medians ranging from 0.2 to 1.2 (Table S5, Figure 2F), though the tracking was significant (upper CI < 1) for few species: Blue Tit, Great Tit, Nuthatch, Pied Flycatcher, and Sedge Warbler (Table S5, Figure 2F).

### Thermal niche tracking across the elevation thermal gradient

Incubation temperatures did not change significantly across elevation for Long-tailed Tit (Table S4, Figure 2), consistent with the thermal niche being quite constant. Significant changes in average fixed window temperature were instead observed across elevation for all species (Table S4, Figure 2G). The observed incubation trend slopes were significantly shallower than average fixed window slopes for 5 species: Blue Tit, Great Tit, Nuthatch, Pied Flycatcher, and Spotted Flycatcher (Table S4, Figure 2G).

The strongest tracking across elevation was observed for Sedge Warbler, followed by Willow Warbler, Wood Warbler, Common Redstart, Nuthatch, Great Tit, Blue Tit, and Pied Flycatcher (Table S5, Figure 2H). In contrast, Spotted Flycatcher exhibited tracking metrics with coefficients and credible intervals above 1 (Table S5, Figure 2H), though these species were identified as exhibiting fitness versus lay date trends consistent with having second broods.

### Cross-species averages for thermal niche tracking

Overall—across both residents and migrants—the tracking metrics indicated partial thermal niche tracking along all four gradients (Table 2). The year tracking had a metric median of 0.50 but its credible intervals slightly overlapped 1 (95% CI: 0.23 to 1.01). The strongest signal was for interannual tracking (0.45, 95% CI: 0.33 to 0.61), followed by latitude (0.65, 95% CI: 0.50 to 0.83) and elevation (0.69, 95% CI: 0.57 to 0.84).

**Table 2:** Thermal niche tracking metric across years, interannual variation, latitudes and elevations on resident and migratory passerines species across the UK for the past 40-50 years. The metrics are calculated as the ratio between observed trends in incubation temperatures and trends in fixed window temperatures. A value of 0 indicates perfect thermal niche tracking, while a value of 1 indicates no tracking. Reported coefficients are the median tracking metric across the posterior for each temporal and spatial dimension. Columns show results based on all species (All) and excluding potentially second-brooded species (Excl.: Common Redstart, Garden Warbler, Spotted Flycatcher, and Willow Warbler). Coefficients and credible intervals (95% CIs) that do not overlap 0 or 1 are highlighted in bold.

|  | All species |  |  |  | Exclusion of second brood species |  |  |  |
| --- | --- | --- | --- | --- | --- | --- | --- | --- |
|  | Resident | Migratory | Resident – Migrant | Overall | Resident | Migratory | Resident – Migrant | Overall |
| Year | 0.37<br>(0.09, 1.18) | 0.61<br>(0.26, 1.49) | -0.24<br>(-1.13, 0.62) | 0.50<br>(0.23, 1.01) | 0.37<br>(0.09, 1.19) | 0.64<br>(0.20, 2.67) | -0.27<br>(-2.29, 0.64) | 0.46<br>(0.18, 1.07) |
| Interannual | **0.52**<br>**(0.34, 0.84)** | **0.40**<br>**(0.28, 0.59)** | 0.11<br>(-0.14, 0.46) | **0.45**<br>**(0.33, 0.61)** | **0.52**<br>**(0.34, 0.85)** | **0.46**<br>**(0.28, 0.72)** | 0.06<br>(-0.26, 0.42) | **0.49**<br>**(0.35, 0.70)** |
| Latitude | **0.50**<br>**(0.34, 0.73)** | **0.77**<br>**(0.59, 0.98)** | -0.26<br>(-0.53, 0.02) | **0.65**<br>**(0.50, 0.83)** | **0.50**<br>**(0.34, 0.73)** | **0.68**<br>**(0.47, 0.96)** | -0.17<br>(-0.50, 0.13) | **0.55**<br>**(0.41, 0.75)** |
| Elevation | **0.56**<br>**(0.42, 0.70)** | 0.88<br>(0.73, 1.05)<sup>1</sup> | **-0.32**<br>**(-0.54, -0.11)<sup>1</sup>** | **0.69**<br>**(0.57, 0.84)<sup>1</sup>** | **0.56**<br>**(0.42, 0.70)** | 0.74<br>(0.57, 1.02)<sup>2</sup> | -0.18<br>(-0.48, 0.04)<sup>2</sup> | **0.62**<br>**(0.50, 0.74)<sup>2</sup>** |

<sup>1</sup> This estimate excludes Garden Warbler and Sedge Warbler due to their lower elevational extent, see methods.
<sup>2</sup> This estimate excludes Sedge Warbler due to their lower elevational extent.

Excluding the species for which we had concerns about second-brooding species yielded similar trends (Table 2).

When we compared the overall tracking metrics across different gradients, we found that on average the focal species tracked their thermal niche more closely across interannual variations than across elevation (significantly smaller tracking metric across interannual variations; Table S6). After excluding the double-brooded species, we found broadly similar patterns but in no case was the difference between tracking metrics significant (i.e., credible intervals for the difference in tracking included zero Table S6).

### Thermal niche tracking comparisons between resident and migratory species

Across the year gradient, resident species had a tracking metric of 0.37 (95% CI: 0.09 to 1.18), compared to 0.61 (95% CI: 0.26 to 1.49) for migratory species (Table 1), but the difference between the groups was not significant (−0.24, 95% CI: -1.13 to 0.62, Table 2), contrary to our hypothesis that resident species are more effective than migratory species at tracking their incubation thermal niche across years.

With respect to interannual variation, resident species had a tracking metric of 0.52 (95% CI: 0.34 to 0.84), compared to 0.40 (95% CI: 0.28 to 0.59) for migratory species (Table 2). The difference between these was not significant (0.11, 95% CI: -0.14 to 0.46), in contrast with our hypothesis that residents would outperform migrants in terms of tracking an incubation thermal niche interannually.

Across latitudes, resident species had a tracking metric of 0.50 (95% CI: 0.34 to 0.73, Table 2), compared to 0.77 (95% CI: 0.59 to 0.98) for migratory species. There was no significant difference between these metrics (−0.26, 95% CI: -0.53 to 0.02), contrary to our hypothesis that migrants would better track thermal niche across latitude.

Across elevations, residents had a tracking metric of 0.56 (95% CI: 0.42 to 0.70), compared to 0.88 (95% CI: 0.73 to 1.05) for migratory species (Table 2). The difference between the metrics was significant (−0.32, 95% CI: -0.54 to -0.11), consistent with resident species more closely tracking an incubation thermal niche across elevations.

After excluding species for which we were concerned about double-brooding, most trends remained qualitatively consistent, except that the difference between resident and migrant species in tracking across elevations was smaller and no longer significant (Table 2).

### Selection on the thermal niche

Three species showed a significant effect of minimum incubation temperature on hatching probability (Table S8, Figure 3). For Blue Tit we estimated a quadratic effect, with hatching probability maximised at intermediate incubation temperatures (Table S8, Figure 3A). For Great Tit and Long-tailed Tit we estimated a negative linear effect, with the hatching probability of Great Tits decreasing by -0.05 logit units (CI = -0.08, -0.01, Table S8, Figure 3C) and Long-tailed Tit by -0.07 logit units (CI = -0.14, -0.01, Table S8, Figure 3E) per 1°C increase in temperature. For all other species, minimum incubation temperatures did not significantly affect hatching probability (Table S8, Figure S16).

**Figure 3:**
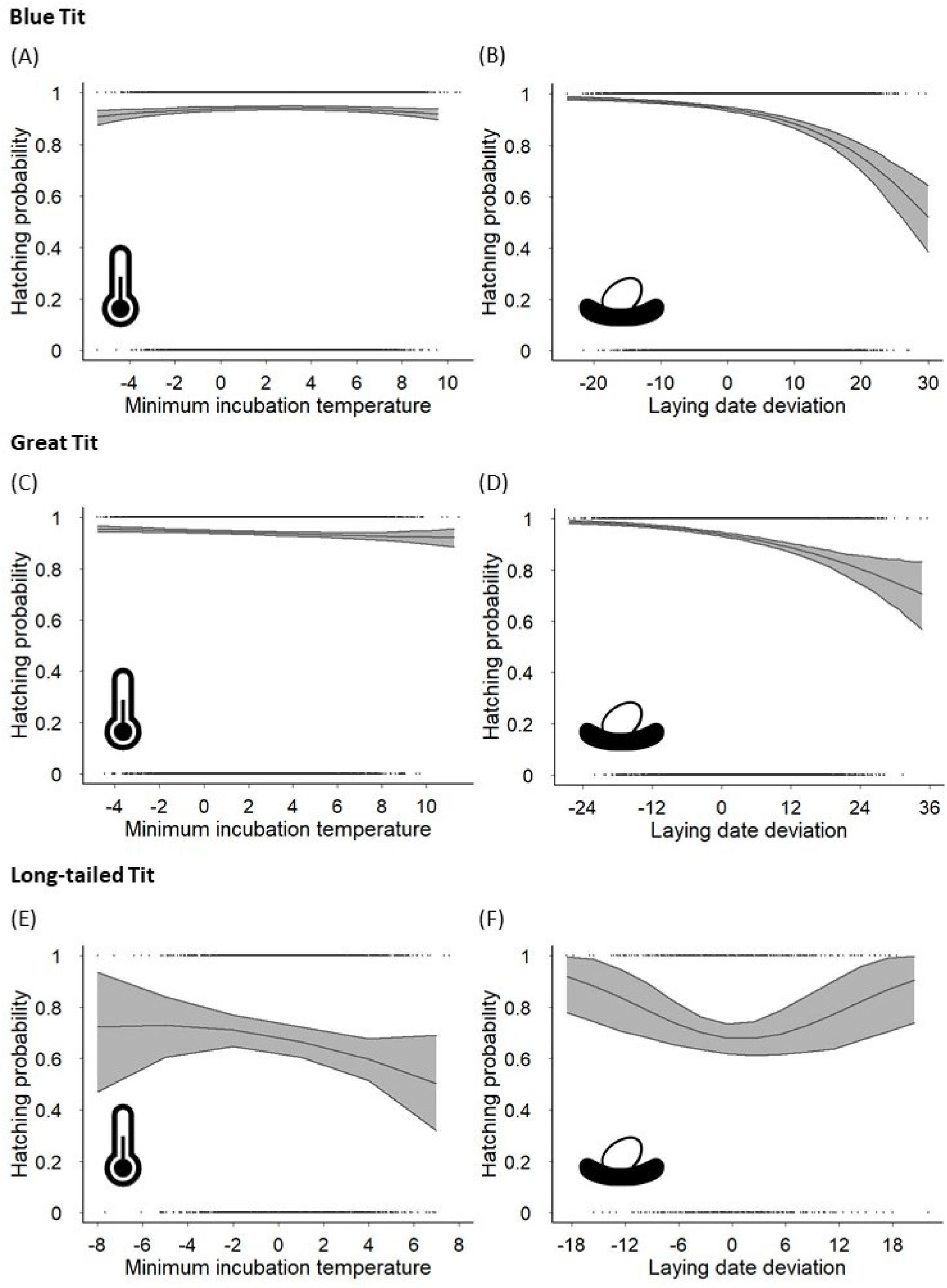
Trends in hatching probability in response to minimum incubation temperature (left column) and laying date deviations (right column) for (A-B) Blue Tit, (C-D) Great Tit, and (E-F) Long-tailed Tit, for the past 40-50 years, across the UK. Hatching probability was measured using a binary metric, with nests containing at least one alive chick at least on day 5 post-hatching classified as successful (nest success = 1) and nests with no alive chicks considered unsuccessful (nest success = 0). The model predictions and 95% credible intervals are reported.

## Discussion

In this study, we tested the extent to which passerine birds track an incubation thermal niche across multiple temporal and spatial gradients in the UK. Most species exhibited partial tracking of the thermal niche across years, interannual variations, latitudes, and elevations, although in many instances we could not reject the null hypothesis of no tracking. Averaging across species we found evidence for significant partial tracking across interannual variation, latitude, and elevation, but not across years. Contrary to our predictions, we found little evidence for tracking ability differing between resident and migratory species across most gradients. The exception was the elevational gradient, where resident species exhibited significantly stronger tracking than migrants, though this difference did not remain after removing species for which we suspected some instances of second brooding. Despite evidence of partial tracking across most gradients, we found weak evidence of selection on the incubation thermal niche for most species. In fact, we only found a significant effect of ambient temperature during incubation on our hatching probability fitness approximation in three resident species, and only for one of these was selection on incubation temperature consistent with our predictions. Overall, we show that conceptualising phenological responses in terms of a species’ thermal niche provides a valuable abiotic yardstick for assessing how organisms are responding to climate change.

Across species, we found no significant evidence of thermal niche tracking across the long-term year gradient, despite moderate overall warming. This result was unexpected given earlier work showing long-term phenological advancements in birds (Charmantier et al., 2008; McLean et al., 2022; Visser et al., 2006, 2015), and recent studies showing how long-term breeding phenology can offset spring warming (López-Idiáquez et al., 2024; Socolar et al., 2017). Our finding rather suggests that across these species phenological advances may not fully compensate for the ambient temperature increases, which is similar to findings in sea turtles (Laloë & Hays, 2023). Whilst we are unable to reject the null hypothesis of no thermal tracking, the median of the tracking metric is consistent with partial tracking. Our finding of very wide credible intervals for this axis, can be attributed to the fact that the average rate of warming over the focal time-period in the UK has been quite small (∼0.03 °C/year, which equals ∼1.5 °C over 50 years) and highly variable across the different fixed windows. Our inability to reject the null hypothesis of no thermal tracking is in contrast with the pronounced thermal tracking reported for Great Tits in Wytham woods over an almost 60-year period (López-Idiáquez et al., 2024). We think this difference may be attributable to the different ways in which heterogeneity in the temperature trend in fixed windows feeds into the calculation; whilst our analysis incorporates this heterogeneity in thermal trend across different stages in the spring, López-Idiáquez et al. (2024) compare the observed thermal change to a single value for the average thermal change expected under the null. Earlier work has suggested that residents tend to be more plastic in their phenological responses to temperature cues than migrants (Samplonius et al., 2018; Usui et al., 2017) whereas there has been some evidence that migrant species may be advancing their breeding phenology through genetic change (Helm et al., 2019; Lonero et al., 2024).

However, we found no evidence for a difference in the tracking ability of migrant and resident species, though the wide credible intervals for our year-tracking metric means that we will have had little power to make this comparison.

Across species, we found evidence of thermal niche tracking across interannual variation, with all 13 species experiencing lower interannual variance in ambient temperature during incubation than expected under the null. This suggests that populations make flexible adjustments in breeding timing, which partially tracks temperature fluctuations, consistent with widespread evidence of temperate passerines exhibiting phenological plasticity to spring temperatures (e.g., Phillimore et al., 2012, 2016). We predicted that the tendency for migratory species to exhibit less thermal sensitivity in their laying dates than resident species (Lehikoinen et al., 2004; Usui et al., 2017) would translate into a weaker ability for interannual tracking, but we found no evidence for this. In fact, the median thermal tracking estimate is lower across migrants than residents, though the difference is not significant. A possible explanation for this unexpected finding is if migrant species tend to have a greater ability to make adjustments in timings in response to temperatures after laying but prior to incubation, as shown for Tree Swallows (*Tachycineta bicolor*) (Ardia et al., 2006).

Across latitude, on average species showed significant but partial thermal niche tracking, maintaining relatively consistent incubation conditions despite the spatial temperature gradient. This suggests that populations of a species breeding at different latitudes experience broadly comparable thermal environments during incubation. Such consistency implies either plastic shifts in timing to match local conditions (Phillimore et al., 2010) or spatial sorting through dispersal or local adaptation (Lamers et al., 2023).

Contrary to our expectations, we found no difference between resident and migratory species in their ability to track their thermal niche across latitudes. Given a tendency for migratory species to have greater natal dispersal propensity (Paradis et al., 1998) we hypothesised they would exhibit stronger spatial tracking of the thermal niche across latitudes. However, we find no significant difference between migrants and residents, though the point estimate is consistent with resident species being better at tracking. We suggest the strong ability of resident species for spatial thermal tracking may be attributable to the high phenotypic plasticity in breeding phenology (Charmantier et al., 2008; Gienapp et al., 2013; Phillimore et al., 2016). Nonetheless, some of the resident species included here have also undergone poleward range shifts (Massimino et al., 2015), raising the possibility that plasticity and spatial redistribution together may maintain thermal stability (Fredston et al., 2025).

On average, birds also showed significant thermal niche tracking across elevation, suggesting that populations can incubate at roughly consistent ambient temperatures from low to high elevations, despite the pronounced thermal gradient. This tracking may arise from compensatory shifts in timing or very local-scale movements. Unexpectedly, resident species tracked a consistent incubation temperature across elevations more closely than migrants, though this pattern was driven mainly by opposite tracking trends in Garden Warbler, Sedge Warbler and Spotted Flycatcher. One potential explanation for this is a greater tendency to have a second brood at lower elevations (Badyaev & Ghalambor, 2001). Tracking metrics exceeding a value of 1 —which we term counter gradient—indicate that observed incubation temperatures have shifted in the opposite direction to that expected for consistent thermal niche maintenance (i.e., this would be consistent with birds breeding earlier at higher, colder elevations than they do at lower, warmer elevations). For a single species, the Spotted Flycatcher, we estimate a thermal tracking value significantly > 1. One possibility is that this pattern reflects an adaptive response to additional environmental conditions and the constraints on timings encountered at different elevations. For instance, birds selecting territories at higher elevations may need to breed earlier to track local resources due to the shorter season at these gradients. Consistent with this explanation, counter-gradient spatial phenological trends—whereby phenology at higher elevations / lower temperatures is earlier than predicted if all populations responded solely through plasticity with respect to spring temperatures—have been widely reported for temperate plants (Tansey et al., 2017) and invertebrates (Gutiérrez & Wilson, 2021; Roy et al., 2015). A further explanation is that migrants may occupy higher elevations in warmer years to avoid potential phenological asynchronies at lower elevations, resulting in earlier breeding at higher elevations (Shutt et al., 2022).

An important consideration with our analyses is the extent to which reliance on 1km interpolated temperature data introduces measurement error and biases estimates. We anticipate such biases are likely to be most problematic for the elevation analyses, specifically where the elevations of grid-cell centroids differ from the ones at the nest locations. One impact of this will be for us to underestimate the true thermal gradient across elevations in fixed windows, and we do observe a lapse rate of –0.3 °C per 100 m that is shallower than the ∼ –0.6 °C per 100 m expected. However, as minimum temperatures will tend to occur during the night time, a better comparison would be with a night time lapse rate, and these are known to be considerably shallower and can sometimes be inverted (Pepin, 2001). The somewhat coarse spatial resolution also limits our ability to detect local-scale behavioural mechanisms that may underpin thermal tracking. For example, Shutt et al. (2022) showed that interannual thermal tracking in migrants can arise from territory selection over tens to hundreds of metres. Incorporating finer-scale temperature and microhabitat data in future work would therefore help resolve how behavioural flexibility contribute to maintaining a consistent thermal niche.

We found limited evidence for selection on incubation temperature, with significant effects detected in only three resident species. This was unexpected, as we anticipated selection on the incubation thermal niche to act as a driver of thermal niche tracking. Our predictions were met only for Blue Tits, which showed higher hatching probability at intermediate incubation temperatures. A similar effect was reported by López-Idiáquez et al. (2024) for Great Tits. However, unlike their study, our use of a binary hatching-success metric may have limited our power to detect stronger selection patterns. Incorporating more informative measures, such as the proportion of eggs hatching or fledgling, would likely provide more nuanced insights into selection. Our inability to identify optimal temperatures for hatching probability may also reflect indirect effects of temperature on fitness. Socolar et al. (2017) highlighted the cascading effects of spring phenology and temperature shifts on breeding behaviours in North American birds, distinguishing indirect—such as synchronization with food resources—from direct thermal influences during incubation. By including both early spring and nesting-season temperatures in their analyses, they reduced the risk of confounding phenological mismatches with direct thermal effects on nesting. Similarly, our focus on incubation temperatures aimed to isolate the direct thermal impacts during this critical stage, distinct from earlier temperature effects on biotic conditions. While parental incubation behaviour likely buffers embryos from moderate temperature fluctuations, extreme cold ambient temperatures could still have detrimental effects. Additionally, ambient temperatures during incubation may influence fitness by affecting prey availability later in the chick-rearing period, potentially impacting nestling growth and survival, although there is so far little evidence that this is a key driver of long-term population changes in UK passerines (Franks et al., 2018).

Separating direct temperature effects on embryo development from indirect effects related to phenological tracking and resource synchronization remains a major challenge. López-Idiáquez et al. (2024) found that over six decades, Great Tits advanced breeding to maintain a consistent thermal niche during nesting, achieving peak reproductive success at intermediate breeding temperatures. Breeding outside these optimal windows led to mismatches with caterpillar abundance, which they interpreted as evidence that thermal tracking is driven more by indirect effects, such as synchronization with food availability, than by direct thermal constraints. Considering this complexity, our results are best interpreted in the context of both direct and indirect effects of temperature on fitness. We also cannot rule out the possibility that the thermal tracking we observe is driven more by indirect effects of temperature, i.e. species tracking the timing of a temperature-sensitive resource, than by direct tracking of temperature. Regardless of whether the impacts of temperature are direct or indirect, we argue that this framing of tracking provides a useful yardstick against which to judge population responses to climate change, that is complementary to other yardsticks such as tracking food resources (Visser & Both, 2005) and optimum breeding timing (Chevin et al., 2015) relative to environmental conditions. This framework can be applied across taxa and environments to assess the extent to which phenological adjustments maintain exposure to suitable temperatures. We suggest that it could be particularly informative to apply this approach to more extreme, more variable and more rapidly warming environments, where the impacts of the thermal niche on fitness may be more pronounced and where limits to phenological plasticity may become more apparent (Gerlich et al., 2025).

In conclusion, we find that passerine populations are partially tracking thermal variation across time and space, suggesting capacity to buffer reproductive stages against climatic variability. The broadly similar tracking strengths across temporal and spatial gradients indicate that phenological plasticity is likely the dominant mechanism enabling these species to track incubation conditions despite changing temperatures. Our study reveals comparable thermal niche tracking abilities between resident and migratory passerines, which does not support our predictions that residents’ greater phenological plasticity may allow them to track temperature better across years, and that migrants’ greater dispersal may allow them to track temperature better across latitudes and elevations. Although selection on the thermal niche appeared weak for most species, this may reflect limitations of our fitness metric. By framing phenological responses through this thermal niche yardstick, our approach offers a pragmatic and widely applicable complement to existing biotic yardsticks.

## Contribution

Conceptualization (equal), Data Curation (lead), Formal analysis (lead), Investigation (lead), Methodology (lead), Project Administration (lead), Visualization (lead), Writing - original draft (lead).

Conceptualization (equal), Funding Acquisition (lead), Supervision (equal), Writing - review & editing (equal)

Conceptualization (equal), Supervision (equal), Writing - review & editing (equal)

Conceptualization (equal), Supervision (equal), Writing - review & editing (equal)

Conceptualization (equal), Investigation (supporting), Methodology (supporting), Supervision (equal), Writing - review & editing (equal)

## Acknowledgments

We gratefully acknowledge the dedicated efforts of the volunteer recorders, whose passion for contributing data to the British Trust for Ornithology (BTO) was essential to this research. We also thank the reviewers for their thoughtful feedback, which helped to improve the quality of the manuscript. Lastly, we are thankful to the editorial team for their consistent support and guidance throughout the publication process.

## References

1. Ardia, D. R., Cooper, C. B., & Dhondt, A. A. (2006). Warm temperatures lead to early onset of incubation, shorter incubation periods and greater hatching asynchrony in tree swallows *Tachycineta bicolor* at the extremes of their range. Journal of Avian Biology, 37(2), 137–142. 10.1111/j.0908-8857.2006.03747.x

2. Badyaev, A. V., & Ghalambor, C. K. (2001). EVOLUTION OF LIFE HISTORIES ALONG ELEVATIONAL GRADIENTS: TRADE-OFF BETWEEN PARENTAL CARE AND FECUNDITY. Ecology, 82(10), 2948–2960. 10.1890/0012-9658(2001)082[2948:EOLHAE]2.0.CO;2

3. Bell, J. R., Botham, M. S., Henrys, P. A., Leech, D. I., Pearce-Higgins, J. W., Shortall, C. R., Brereton, T. M., Pickup, J., & Thackeray, S. J. (2019). Spatial and habitat variation in aphid, butterfly, moth and bird phenologies over the last half century. Global Change Biology, 25(6), 1982–1994. 10.1111/gcb.14592

4. Bellard, C., Bertelsmeier, C., Leadley, P., Thuiller, W., & Courchamp, F. (2012). Impacts of climate change on the future of biodiversity. Ecology Letters, 15(4), 365–377. 10.1111/j.1461-0248.2011.01736.x

5. Bonnet-Lebrun, A., Somveille, M., Rodrigues, A. S. L., & Manica, A. (2021). Exploring intraspecific variation in migratory destinations to investigate the drivers of migration. Oikos, 130(2), 187–196. 10.1111/oik.07689

6. Both, C., Bouwhuis, S., Lessells, C. M., & Visser, M. E. (2006). Climate change and population declines in a long-distance migratory bird. Nature, 441(7089), 81–83. 10.1038/nature04539

7. Both, C., & Visser, M. E. (2001). Adjustment to climate change is constrained by arrival date in a long-distance migrant bird. Nature, 411(6835), 296–298. 10.1038/35077063

8. BTO. (2025). BirdFacts: Profiles of birds occurring in the United Kingdom. https://www.bto.org/birdfacts

9. Burgess, M. D., Smith, K. W., Evans, K. L., Leech, D., Pearce-Higgins, J. W., Branston, C. J., Briggs, K., Clark, J. R., Du Feu, C. R., Lewthwaite, K., Nager, R. G., Sheldon, B. C., Smith, J. A., Whytock, R. C., Willis, S. G., & Phillimore, A. B. (2018). Tritrophic phenological match–mismatch in space and time. Nature Ecology & Evolution, 2(6), 970–975. 10.1038/s41559-018-0543-1

10. Charmantier, A., McCleery, R. H., Cole, L. R., Perrins, C., Kruuk, L. E. B., & Sheldon, B. C. (2008). Adaptive Phenotypic Plasticity in Response to Climate Change in a Wild Bird Population. Science, 320(5877), 800–803. 10.1126/science.1157174

11. Chen, I.-C., Hill, J. K., Ohlemüller, R., Roy, D. B., & Thomas, C. D. (2011). Rapid Range Shifts of Species Associated with High Levels of Climate Warming. Science, 333(6045), 1024–1026. 10.1126/science.1206432

12. Chevin, L.-M., Visser, M. E., & Tufto, J. (2015). Estimating the variation, autocorrelation, and environmental sensitivity of phenotypic selection: ESTIMATING FLUCTUATING SELECTION. Evolution, 69(9), 2319–2332. 10.1111/evo.12741

13. Cohen, J. M., Lajeunesse, M. J., & Rohr, J. R. (2018). A global synthesis of animal phenological responses to climate change. Nature Climate Change, 8(3), 224–228. 10.1038/s41558-018-0067-3

14. Crick, H. Q. P., Baillie, S. R., & Leech, D. I. (2003). The UK Nest Record Scheme: Its value for science and conservation. Bird Study, 50(3), 254–270. 10.1080/00063650309461318

15. Dunn, E. (1977). Predation by Weasels (Mustela nivalis) on Breeding Tits (Parus Spp.) in Relation to the Density of Tits and Rodents. The Journal of Animal Ecology, 46(2), 633. 10.2307/3835

16. DuRant, S. E., Carter, A. W., Denver, R. J., Hepp, G. R., & Hopkins, W. A. (2014). Are thyroid hormones mediators of incubation temperature-induced phenotypes in birds? Biology Letters, 10(1), 20130950. 10.1098/rsbl.2013.0950

17. DuRant, S. E., Hepp, G. R., Moore, I. T., Hopkins, B. C., & Hopkins, W. A. (2010). Slight differences in incubation temperature affect early growth and stress endocrinology of wood duck (*Aix sponsa*) ducklings. Journal of Experimental Biology, 213(1), 45–51. 10.1242/jeb.034488

18. DuRant, S. E., Hopkins, W. A., Carter, A. W., Stachowiak, C. M., & Hepp, G. R. (2013). Incubation Conditions Are More Important in Determining Early Thermoregulatory Ability than Posthatch Resource Conditions in a Precocial Bird. Physiological and Biochemical Zoology, 86(4), 410–420. 10.1086/671128

19. DuRant, S. E., Hopkins, W. A., Hepp, G. R., & Walters, J. R. (2013). Ecological, evolutionary, and conservation implications of incubation temperature-dependent phenotypes in birds. Biological Reviews, 88(2), 499–509. 10.1111/brv.12015

20. Finch, T., Pearce-Higgins, J. W., Leech, D. I., & Evans, K. L. (2014). Carry-over effects from passage regions are more important than breeding climate in determining the breeding phenology and performance of three avian migrants of conservation concern. Biodiversity and Conservation, 23(10), 2427–2444. 10.1007/s10531-014-0731-5

21. Franks, S. E., Pearce-Higgins, J. W., Atkinson, S., Bell, J. R., Botham, M. S., Brereton, T. M., Harrington, R., & Leech, D. I. (2018). The sensitivity of breeding songbirds to changes in seasonal timing is linked to population change but cannot be directly attributed to the effects of trophic asynchrony on productivity. Global Change Biology, 24(3), 957–971. 10.1111/gcb.13960

22. Fredston, A. L., Tingley, M. W., Neate-Clegg, M. H. C., Evans, L. J., Antão, L. H., Ban, N. C., Chen, I.-C., Chen, Y.-W., Comte, L., Edwards, D. P., Evengard, B., Fadrique, B., Falkeis, S. H., Guralnick, R., Klinges, D. H., Lembrechts, J. J., Lenoir, J., Palacios-Abrantes, J., Pauchard, A., … Scheffers, B. R. (2025). Reimagining species on the move across space and time. Trends in Ecology & Evolution, 40(7), 629–638. 10.1016/j.tree.2025.03.015

23. Gerlich, H. S., Holmstrup, M., Schmidt, N. M., Phillimore, A. B., & Høye, T. T. (2025). Keeping up with climate change: Have Arctic arthropods reached their phenological limits? Proceedings of the Royal Society B: Biological Sciences, 292(2051), 20250350. 10.1098/rspb.2025.0350

24. Gienapp, P., Lof, M., Reed, T. E., McNamara, J., Verhulst, S., & Visser, M. E. (2013). Predicting demographically sustainable rates of adaptation: Can great tit breeding time keep pace with climate change? Philosophical Transactions of the Royal Society B: Biological Sciences, 368(1610), 20120289. 10.1098/rstb.2012.0289

25. Gómez, C., Tenorio, E. A., Montoya, P., & Cadena, C. D. (2016). Niche-tracking migrants and niche-switching residents: Evolution of climatic niches in New World warblers (Parulidae). Proceedings of the Royal Society B: Biological Sciences, 283(1824), 20152458. 10.1098/rspb.2015.2458

26. Gutiérrez, D., & Wilson, R. J. (2021). Intra- and interspecific variation in the responses of insect phenology to climate. Journal of Animal Ecology, 90(1), 248–259. 10.1111/1365-2656.13348

27. Gvoždík, L. (2018). Just what is the thermal niche? Oikos, 127(12), 1701–1710. 10.1111/oik.05563

28. Hadfield, J. D. (2010). MCMC Methods for Multi-Response Generalized Linear Mixed Models: The **MCMCglmm** *R* Package. Journal of Statistical Software, 33(2). 10.18637/jss.v033.i02

29. Helm, B., Van Doren, B. M., Hoffmann, D., & Hoffmann, U. (2019). Evolutionary Response to Climate Change in Migratory Pied Flycatchers. Current Biology, 29(21), 3714–3719.e4. 10.1016/j.cub.2019.08.072

30. Hepp, G. R., Kennamer, R. A., & Johnson, M. H. (2006). Maternal effects in Wood Ducks: Incubation temperature influences incubation period and neonate phenotype. Functional Ecology, 20(2), 308–314. 10.1111/j.1365-2435.2006.01108.x

31. Hollister, J. W. (2023). Elevatr: Access Elevation Data from Various APIs. R package version 0.99.0. [Computer software]. https://CRAN.R-project.org/package=elevatr/

32. Hopkins, A. D. (1920). THE BIOCLIMATIC LAW^1^. Monthly Weather Review, 48(6), 355–355. 10.1175/1520-0493(1920)48<355a:TBL>2.0.CO;2

33. Hulme, M. (2001). Climatic perspectives on Sahelian desiccation: 1973}1998. Global Environmental Change.

34. IPCC. (2007). Climate Change 2007: The Physical Science Basis. Contribution of Working Group I to the Fourth Assessment Report of the Intergovernmental Panel on Climate Change [Solomon, S., D. Qin, M. Manning, Z. Chen, M. Marquis, K.B. Averyt, M. Tignor and H.L. Miller (eds.)]. Cambridge University Press.

35. Laloë, J.-O., & Hays, G. C. (2023). Can a present-day thermal niche be preserved in a warming climate by a shift in phenology? A case study with sea turtles. Royal Society Open Science, 10(2), 221002. 10.1098/rsos.221002

36. Lamers, K. P., Nilsson, J.-Å., Nicolaus, M., & Both, C. (2023). Adaptation to climate change through dispersal and inherited timing in an avian migrant. Nature Ecology & Evolution, 7(11), 1869–1877. 10.1038/s41559-023-02191-w

37. Lehikoinen, E., Sparks, T. H., & Zalakevicius, M. (2004). Arrival and Departure Dates. In Advances in Ecological Research (Vol. 35, pp. 1–31). Elsevier. 10.1016/S0065-2504(04)35001-4

38. Linden, M., & Møller, A. P. (1989). Cost of reproduction and covariation of life history traits in birds. Trends in Ecology & Evolution, 4(12), 367–371. 10.1016/0169-5347(89)90101-8

39. Lonero, I., Eddowes, M. J., Burgess, M. D., Pearce-Higgins, J. W., & Phillimore, A. B. (2024). Temperature sensitivity of breeding phenology and reproductive output of the Common Redstart (*Phoenicurus phoenicurus*). Ibis, ibi.13376. 10.1111/ibi.13376

40. López-Idiáquez, D., Cole, E. F., Regan, C. E., & Sheldon, B. C. (2024). Optimal thermal niche-tracking buffers wild great tits against climate change. 10.1101/2024.07.18.603942

41. Low, M., & Pärt, T. (2009). Patterns of mortality for each life-history stage in a population of the endangered New Zealand stitchbird. Journal of Animal Ecology, 78(4), 761–771. 10.1111/j.1365-2656.2009.01543.x

42. Lozano Ruiz, C. (2022). Sgo: Simple Geographical Operations (with OSGB36). R package version 0.9.2. [Computer software]. https://CRAN.R-project.org/package=sgo

43. Macphie, K. H., Samplonius, J. M., Pick, J. L., Hadfield, J. D., & Phillimore, A. B. (2023). Modelling thermal sensitivity in the full phenological distribution: A new approach applied to the spring arboreal caterpillar peak. Functional Ecology, 37(12), 3015–3026. 10.1111/1365-2435.14436

44. Magnuson, J. J., Crowder, L. B., & Medvick, P. A. (1979). Temperature as an Ecological Resource. American Zoologist, 19(1), 331–343. 10.1093/icb/19.1.331

45. Martin, T. E., Auer, S. K., Bassar, R. D., Niklison, A. M., & Lloyd, P. (2007). GEOGRAPHIC VARIATION IN AVIAN INCUBATION PERIODS AND PARENTAL INFLUENCES ON EMBRYONIC TEMPERATURE. Evolution, 61(11), 2558–2569. 10.1111/j.1558-5646.2007.00204.x

46. Massicotte, P., & South, A. (2023). Rnaturalearth: World Map Data from Natural Earth. R package version 1.0.1. [Computer software]. https://CRAN.R-project.org/package=rnaturalearth

47. Massimino, D., Johnston, A., & Pearce-Higgins, J. W. (2015). The geographical range of British birds expands during 15 years of warming. Bird Study, 62(4), 523–534. 10.1080/00063657.2015.1089835

48. McCowan, L. S. C., & Griffith, S. C. (2021). Baked eggs: Catastrophic heatwave-induced reproductive failure in the desert-adapted Zebra Finch (*Taeniopygia guttata*). Ibis, 163(4), 1207–1216. 10.1111/ibi.12958

49. McLean, N., Kruuk, L. E. B., Van Der Jeugd, H. P., Leech, D., Van Turnhout, C. A. M., & Van De Pol, M. (2022). Warming temperatures drive at least half of the magnitude of long-term trait changes in European birds. Proceedings of the National Academy of Sciences, 119(10), e2105416119. 10.1073/pnas.2105416119

50. Monroe, A. P., Hallinger, K. K., Brasso, R. L., & Cristol, D. A. (2008). OCCURRENCE AND IMPLICATIONS OF DOUBLE BROODING IN A SOUTHERN POPULATION OF TREE SWALLOWS. The Condor, 110(2), 382–386. 10.1525/cond.2008.8341

51. Müller, M., Pasinelli, G., Schiegg, K., Spaar, R., & Jenni, L. (2005). Ecological and social effects on reproduction and local recruitment in the red-backed shrike. Oecologia, 143(1), 37–50. 10.1007/s00442-004-1770-5

52. Neate-Clegg, M. H. C., Tonelli, B. A., & Tingley, M. W. (2024). Advances in breeding phenology outpace latitudinal and elevational shifts for North American birds tracking temperature. Nature Ecology & Evolution, 8(11), 2027–2036. 10.1038/s41559-024-02536-z

53. Nilsson, S. G. (1984). The Evolution of Nest-Site Selection among Hole-Nesting Birds: The Importance of Nest Predation and Competition. Ornis Scandinavica, 15(3), 167. 10.2307/3675958

54. Ockendon, N., Leech, D., & Pearce-Higgins, J. W. (2013). Climatic effects on breeding grounds are more important drivers of breeding phenology in migrant birds than carry-over effects from wintering grounds. Biology Letters, 9(6), 20130669. 10.1098/rsbl.2013.0669

55. Paradis, E., Baillie, S. R., Sutherland, W. J., & Gregory, R. D. (1998). Patterns of natal and breeding dispersal in birds. Journal of Animal Ecology, 67(4), 518–536. 10.1046/j.1365-2656.1998.00215.x

56. Parmesan, C. (2006). Ecological and Evolutionary Responses to Recent Climate Change. Annual Review of Ecology, Evolution, and Systematics, 37(1), 637–669. 10.1146/annurev.ecolsys.37.091305.110100

57. Parmesan, C., & Yohe, G. (2003). A globally coherent fingerprint of climate change impacts across natural systems. Nature, 421(6918), 37–42. 10.1038/nature01286

58. Pearce-Higgins, J. W., Eglington, S. M., Martay, B., & Chamberlain, D. E. (2015). Drivers of climate change impacts on bird communities. Journal of Animal Ecology, 84(4), 943–954. 10.1111/1365-2656.12364

59. Pepin, N. (2001). Lapse rate changes in northern England. Theoretical and Applied Climatology, 68(1–2), 1–16. 10.1007/s007040170049

60. Perry, M., Hollis, D., & Elms, M. (2009). The Generation of Daily Gridded Datasets of Temperature and Rainfall for the UK.

61. Phillimore, A. B., Hadfield, J. D., Jones, O. R., & Smithers, R. J. (2010). Differences in spawning date between populations of common frog reveal local adaptation. Proceedings of the National Academy of Sciences, 107(18), 8292–8297. 10.1073/pnas.0913792107

62. Phillimore, A. B., Leech, D. I., Pearce-Higgins, J. W., & Hadfield, J. D. (2016). Passerines may be sufficiently plastic to track temperature-mediated shifts in optimum lay date. Global Change Biology, 22(10), 3259–3272. 10.1111/gcb.13302

63. Phillimore, A. B., Stålhandske, S., Smithers, R. J., & Bernard, R. (2012). Dissecting the Contributions of Plasticity and Local Adaptation to the Phenology of a Butterfly and Its Host Plants. The American Naturalist, 180(5), 655–670. 10.1086/667893

64. Pörtner, H. O., & Farrell, A. P. (2008). Physiology and Climate Change. Science, 322(5902), 690–692. 10.1126/science.1163156

65. R Core Team. (2025). R: a Language and Environment for Statistical Computing. [Computer software]. R Foundation for Statistical Computing. https://www.R-project.org/

66. Radchuk, V., Reed, T., Teplitsky, C., Van De Pol, M., Charmantier, A., Hassall, C., Adamík, P., Adriaensen, F., Ahola, M. P., Arcese, P., Miguel Avilés, J., Balbontin, J., Berg, K. S., Borras, A., Burthe, S., Clobert, J., Dehnhard, N., De Lope, F., Dhondt, A. A., … Kramer-Schadt, S. (2019). Adaptive responses of animals to climate change are most likely insufficient. Nature Communications, 10(1), 3109. 10.1038/s41467-019-10924-4

67. Reid, J. M., Monaghan, P., & Ruxton, G. D. (2000). The consequences of clutch size for incubation conditions and hatching success in starlings. Functional Ecology, 14(5), 560–565. 10.1046/j.1365-2435.2000.t01-1-00446.x

68. Roslin, T., Antão, L., Hällfors, M., Meyke, E., Lo, C., Tikhonov, G., Delgado, M. D. M., Gurarie, E., Abadonova, M., Abduraimov, O., Adrianova, O., Akimova, T., Akkiev, M., Ananin, A., Andreeva, E., Andriychuk, N., Antipin, M., Arzamascev, K., Babina, S., … Ovaskainen, O. (2021). Phenological shifts of abiotic events, producers and consumers across a continent. Nature Climate Change, 11(3), 241–248. 10.1038/s41558-020-00967-7

69. Roy, D. B., Oliver, T. H., Botham, M. S., Beckmann, B., Brereton, T., Dennis, R. L. H., Harrower, C., Phillimore, A. B., & Thomas, J. A. (2015). Similarities in butterfly emergence dates among populations suggest local adaptation to climate. Global Change Biology, 21(9), 3313–3322. 10.1111/gcb.12920

70. Saino, N., Rubolini, D., Jonzén, N., Ergon, T., Montemaggiori, A., Stenseth, N., & Spina, F. (2007). Temperature and rainfall anomalies in Africa predict timing of spring migration in trans-Saharan migratory birds. Climate Research, 35, 123–134. 10.3354/cr00719

71. Salaberria, C., Celis, P., López-Rull, I., & Gil, D. (2014). Effects of temperature and nest heat exposure on nestling growth, dehydration and survival in a Mediterranean hole-nesting passerine. Ibis, 156(2), 265–275. 10.1111/ibi.12121

72. Samplonius, J. M., Bartošová, L., Burgess, M. D., Bushuev, A. V., Eeva, T., Ivankina, E. V., Kerimov, A. B., Krams, I., Laaksonen, T., Mägi, M., Mänd, R., Potti, J., Török, J., Trnka, M., Visser, M. E., Zang, H., & Both, C. (2018). Phenological sensitivity to climate change is higher in resident than in migrant bird populations among European cavity breeders. Global Change Biology, 24(8), 3780–3790. 10.1111/gcb.14160

73. Selonen, V., Helle, S., Laaksonen, T., Ahola, M. P., Lehikoinen, E., & Eeva, T. (2021). Identifying the paths of climate effects on population dynamics: Dynamic and multilevel structural equation model around the annual cycle. Oecologia, 195(2), 525–538. 10.1007/s00442-020-04817-3

74. Shipley, J. R., Twining, C. W., Taff, C. C., Vitousek, M. N., Flack, A., & Winkler, D. W. (2020). Birds advancing lay dates with warming springs face greater risk of chick mortality. Proceedings of the National Academy of Sciences, 117(41), 25590–25594. 10.1073/pnas.2009864117

75. Shutt, J. D., Bell, S. C., Bell, F., Castello, J., El Harouchi, M., & Burgess, M. D. (2022). Territory-level temperature influences breeding phenology and reproductive output in three forest passerine birds. Oikos, 2022(8), e09171. 10.1111/oik.09171

76. Socolar, J. B., Epanchin, P. N., Beissinger, S. R., & Tingley, M. W. (2017). Phenological shifts conserve thermal niches in North American birds and reshape expectations for climate-driven range shifts. Proceedings of the National Academy of Sciences, 114(49), 12976–12981. 10.1073/pnas.1705897114

77. Strode, P. K. (2003). Implications of climate change for North American wood warblers (Parulidae). Global Change Biology, 9(8), 1137–1144. 10.1046/j.1365-2486.2003.00664.x

78. Tansey, C. J., Hadfield, J. D., & Phillimore, A. B. (2017). Estimating the ability of plants to plastically track temperature-mediated shifts in the spring phenological optimum. Global Change Biology, 23(8), 3321–3334. 10.1111/gcb.13624

79. Tarof, S. A., Kramer, P. M., Hill, J. R., Tautin, J., & Stutchbury, B. J. M. (2011). Brood size and late breeding are negatively related to juvenile survival in a Neotropical migratory songbird. The Auk, 128(4), 716–725. 10.1525/auk.2011.11087

80. Thackeray, S. J., Henrys, P. A., Hemming, D., Bell, J. R., Botham, M. S., Burthe, S., Helaouet, P., Johns, D. G., Jones, I. D., Leech, D. I., Mackay, E. B., Massimino, D., Atkinson, S., Bacon, P. J., Brereton, T. M., Carvalho, L., Clutton-Brock, T. H., Duck, C., Edwards, M., … Wanless, S. (2016). Phenological sensitivity to climate across taxa and trophic levels. Nature, 535(7611), 241–245. 10.1038/nature18608

81. Usui, T., Butchart, S. H. M., & Phillimore, A. B. (2017). Temporal shifts and temperature sensitivity of avian spring migratory phenology: A phylogenetic meta-analysis. Journal of Animal Ecology, 86(2), 250–261. 10.1111/1365-2656.12612

82. Vedder, O., Bouwhuis, S., & Sheldon, B. C. (2013). Quantitative Assessment of the Importance of Phenotypic Plasticity in Adaptation to Climate Change in Wild Bird Populations. PLoS Biology, 11(7), e1001605. 10.1371/journal.pbio.1001605

83. Verhulst, S., & Nilsson, J.-Å. (2008). The timing of birds’ breeding seasons: A review of experiments that manipulated timing of breeding. Philosophical Transactions of the Royal Society B: Biological Sciences, 363(1490), 399–410. 10.1098/rstb.2007.2146

84. Verhulst, S., & Tinbergen, J. M. (1991). Experimental Evidence for a Causal Relationship between Timing and Success of Reproduction in the Great Tit Parus m. Major. The Journal of Animal Ecology, 60(1), 269. 10.2307/5459

85. Visser, M. E., & Both, C. (2005). Shifts in phenology due to global climate change: The need for a yardstick. Proceedings of the Royal Society B: Biological Sciences, 272(1581), 2561–2569. 10.1098/rspb.2005.3356

86. Visser, M. E., Both, C., & Lambrechts, M. M. (2004). Global Climate Change Leads to Mistimed Avian Reproduction. In Advances in Ecological Research (Vol. 35, pp. 89–110). Elsevier. 10.1016/S0065-2504(04)35005-1

87. Visser, M. E., Gienapp, P., Husby, A., Morrisey, M., De La Hera, I., Pulido, F., & Both, C. (2015). Effects of Spring Temperatures on the Strength of Selection on Timing of Reproduction in a Long-Distance Migratory Bird. PLOS Biology, 13(4), e1002120. 10.1371/journal.pbio.1002120

88. Visser, M. E., Holleman, L. J. M., & Gienapp, P. (2006). Shifts in caterpillar biomass phenology due to climate change and its impact on the breeding biology of an insectivorous bird. Oecologia, 147(1), 164–172. 10.1007/s00442-005-0299-6

89. Webb, D. R. (1987). Thermal Tolerance of Avian Embryos: A Review. The Condor, 89(4), 874. 10.2307/1368537

90. Williams, J. B. (1996). Energetics of Avian Incubation. In C. Carey (Ed.), Avian Energetics and Nutritional Ecology (pp. 375–415). Springer US. 10.1007/978-1-4613-0425-8_11

91. Winkler, D. W., Luo, M. K., & Rakhimberdiev, E. (2013). Temperature effects on food supply and chick mortality in tree swallows (Tachycineta bicolor). Oecologia, 173(1), 129–138. 10.1007/s00442-013-2605-z

