## Supplementary Material for "Thermal niche tracking in thirteen British temperate passerines"

**Table S8:** Incubation temperature effects on hatching probability .................... 18
**Figure S16:** Plot of of incubation temperature and laying date deviations effects on hatching probability .................... 22
**References** .................... **23**

### Methods

#### *Chick age*

We used the chick age variable to measure the fitness metric of hatching probability, with nests containing at least one alive chick at least on day 5 post-hatching classified as successful (nest success = 1) and nests with no alive chicks considered unsuccessful (nest success = 0).

We measured chick age as the difference in days between the brood's first hatching date (midpoint between minimum and maximum estimates of first hatching dates) and the visit date when the chicks were observed.

#### *Method used to test temperature trends during the birds' incubation period and during fixed time windows*

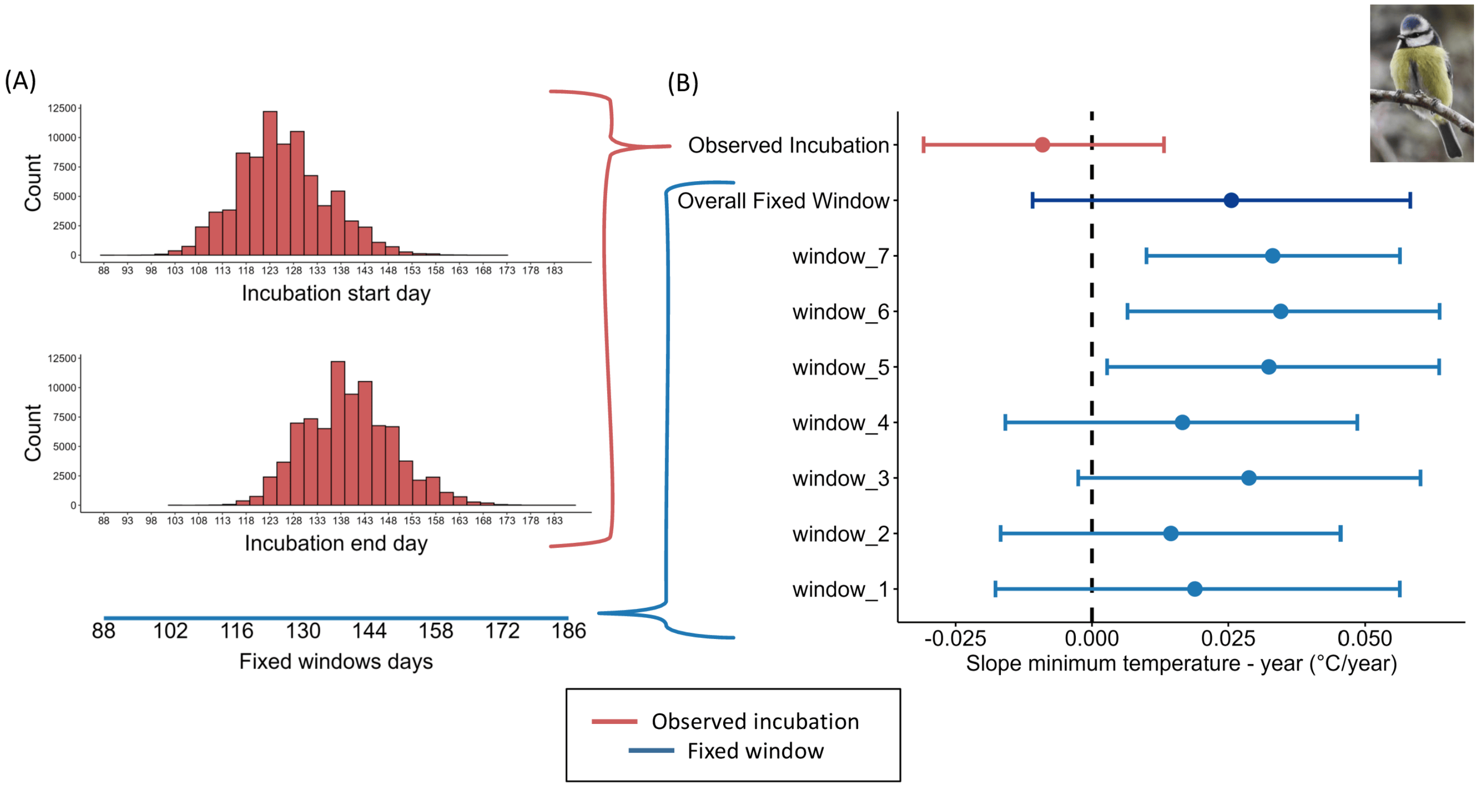

**Figure S1:** Schematic of the method used to test temperature trends during the birds' incubation period and during fixed time windows, shown here as an example for Blue Tits and trends along the long-term temporal gradient. The method consists of two steps: (A) selecting the time windows for analysing null temperature trends and (B) performing the trend analysis within those windows. (A) Fixed time windows (light blue) were chosen based on the species-specific incubation start and end dates (red), ensuring alignment with the incubation period experienced by individual birds of that species. (B) Temperature trends were then compared for one of the considered dimensions (in this example, the temporal gradient) during both the birds' actual incubation period (red) and the corresponding null fixed time windows (light blue). Finally, an overall null temperature trend was calculated across all fixed time windows (dark blue) and this was used for statistical comparisons and in calculation of the thermal niche tracking metric.

#### *Counter-gradient thermal niche tracking*

For each species, the thermal niche tracking metric, referred to as the tracking metric, was calculated as the ratio between the observed trends (median of the posterior slopes and 95% CIs) in incubation temperatures and the trends in fixed window temperatures (median of the posterior slopes and 95% CIs). A tracking metric of 0 would then indicate perfect thermal niche tracking, while a value of 1 would indicate no tracking. Values in the range 0-1 would correspond to partial tracking and values >1 would correspond to what we define as counter-gradient tracking (Figure S2B).

If a species experiences consistent temperatures over time and space during a specific stage of its life cycle, it indicates full thermal niche tracking, achieved through phenological adjustments along environmental gradients (Figure S2A). Conversely, if a species does not experience consistent temperatures but instead encounters more extreme temperatures than expected under the null hypothesis, we describe this as counter-gradient thermal niche tracking (Figure S2B). This pattern suggests that incubation temperatures shift in the opposite direction to what is expected for full thermal niche tracking—for instance, with earlier breeding occurring in later/warmer years, at lower/warmer latitudes, and at lower/warmer elevations, and vice versa.

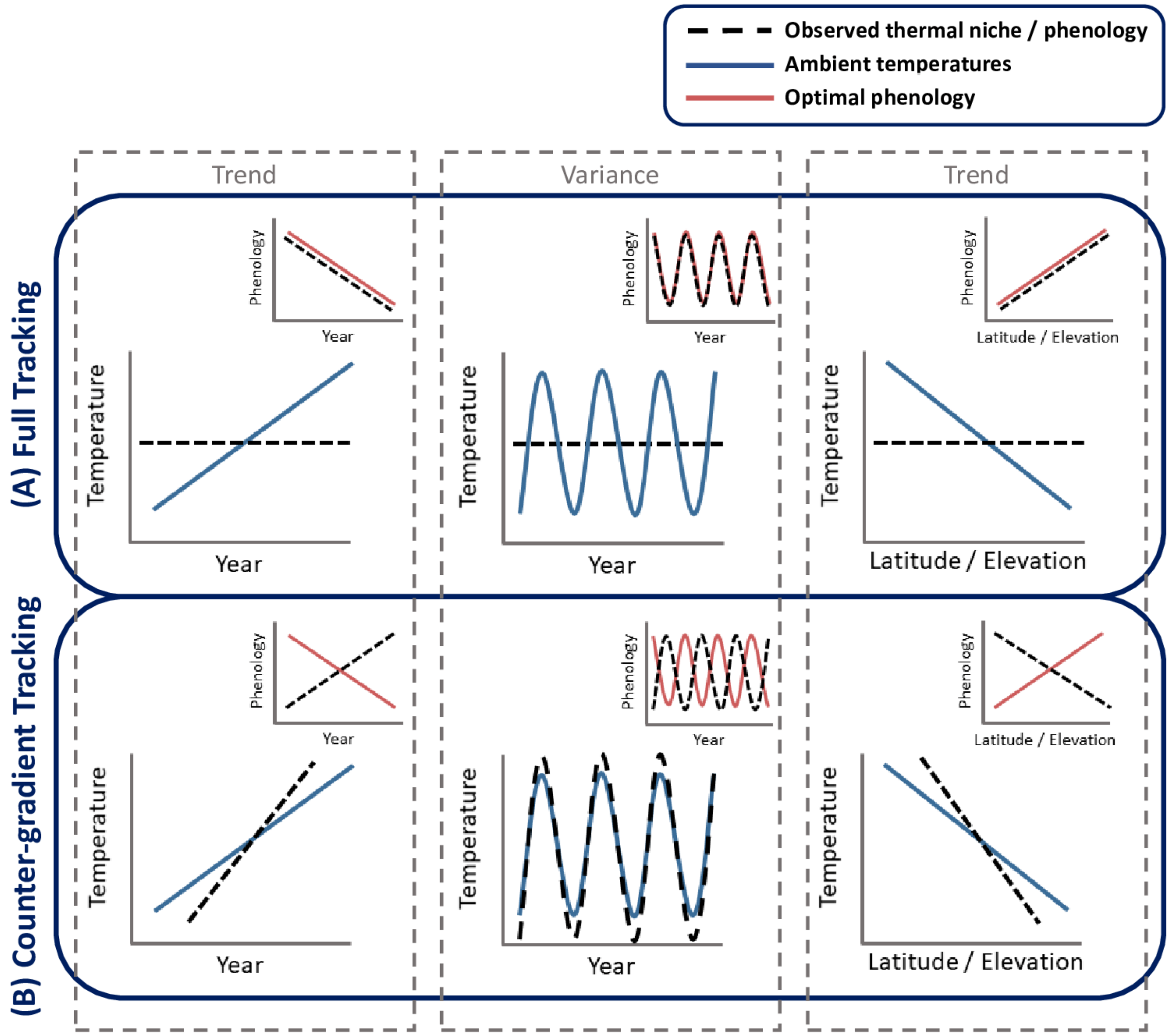

**Figure S2:** Full thermal niche tracking and counter-gradient thermal niche tracking. The schematic illustrates potential scenarios for thermal niche tracking across time (year) and space (latitude, elevation): (A) Full thermal niche tracking, (B) Counter-gradient thermal niche tracking, (C) No thermal niche tracking. Each scenario consists of two types of panels: temperature panels (large panels) and phenology panels (inset panels). The temperature panels display temporal (trend or interannual) and spatial (latitude or elevation trends) temperature. The blue line represents ambient temperatures, and the dashed black line shows temperatures relevant to the species' thermal niche (the temperatures experienced by the species). The phenology panels illustrate temporal and spatial shifts in phenology. The red line represents the optimal phenology for full thermal niche maintenance, while the dashed black line shows realised phenology. This figure helps compare the extent to which realised phenology aligns with the optimal phenology.

### Model Q-Q-Plots

To assess model performance, we examined normal Q-Q plots for the univariate MCMCglmm species models on observed incubation temperatures and fixed (null) windows temperatures. To generate the expectation, we simulated under the model 1000 times and used this output to obtain average quantiles.

The Q-Q plots for the Blue Tit (Fig S3), Chaffinch (Fig S4), Great Tit (Fig S5), Long-tailed Tit (Fig S6), Nuthatch (Fig S7), Common Redstart (Fig S8), Garden Warbler (Fig S9), Pied Flycatcher (Fig S10), Spotted Flycatcher (Fig S12), and Willow Warbler (Fig S14) are quite well-behaved. The Q-Q plots for Sedge Warbler (Fig S11), Whinchat (Fig S13) and Wood Warbler (Fig S15) reveal that the fixed (null) models tend to miss extreme cold events and extreme warm anomalies. The models of Sedge Warbler and Whinchat tend to predict lower cold temperatures and higher warm temperatures than expected, while the models of Wood Warbler tend to either under- or over-predict lower and higher temperatures, depending on the window.

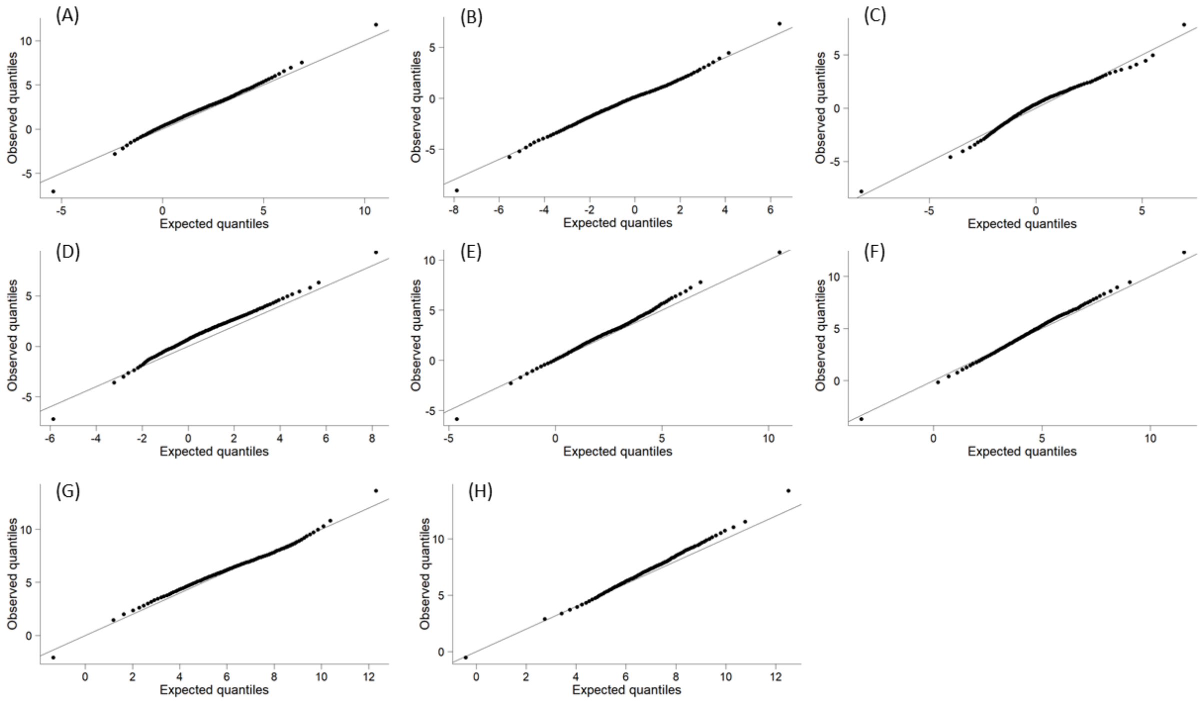

**Figure S3:** Q-Q plots for the model examining temporal and geographic trends in the minimum temperature during the observed incubation period for the Blue Tit (A), and for the models examining temporal and geographic trends in the minimum temperature during fixed (null) windows for the Blue Tit (B – H).

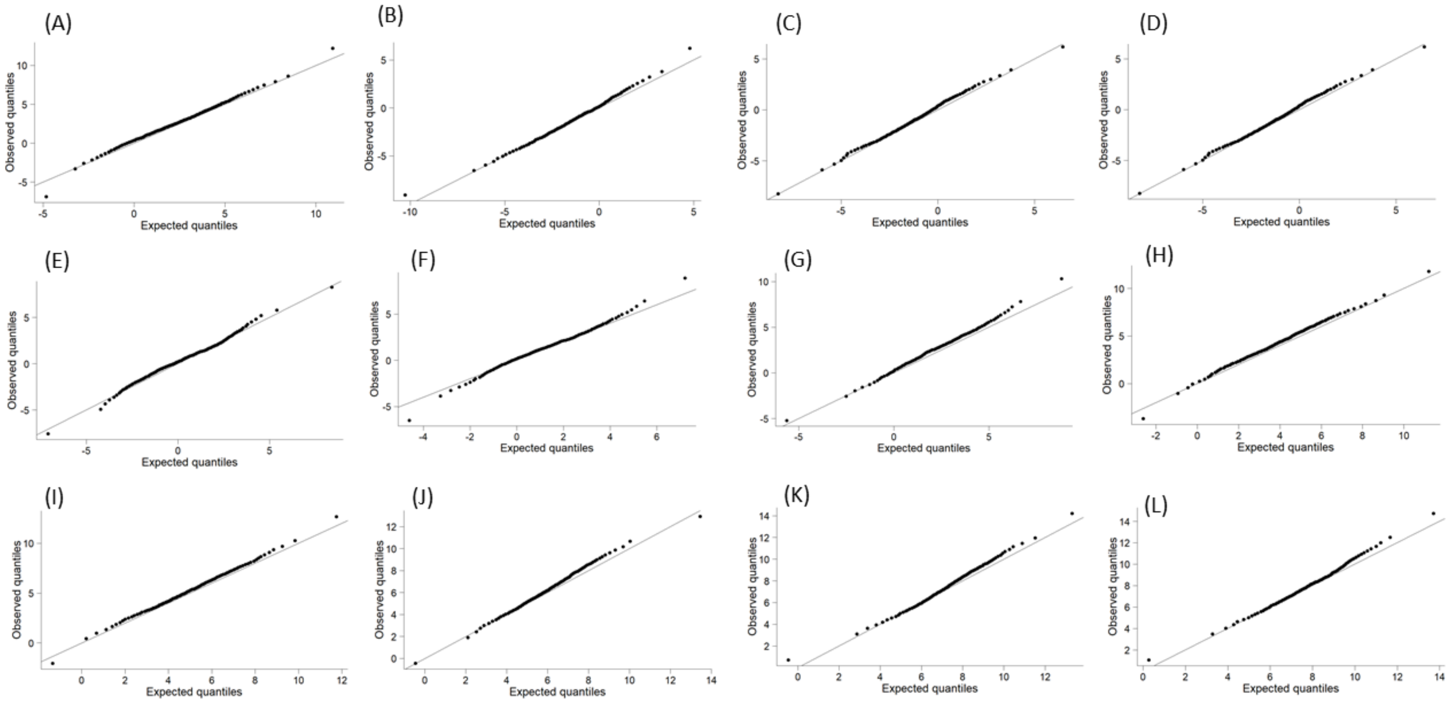

**Figure S4:** Q-Q plots for the model examining temporal and geographic trends in the minimum temperature during the observed incubation period for the Chaffinch (A), and for the models examining temporal and geographic trends in the minimum temperature during fixed (null) windows for the Chaffinch (B – L).

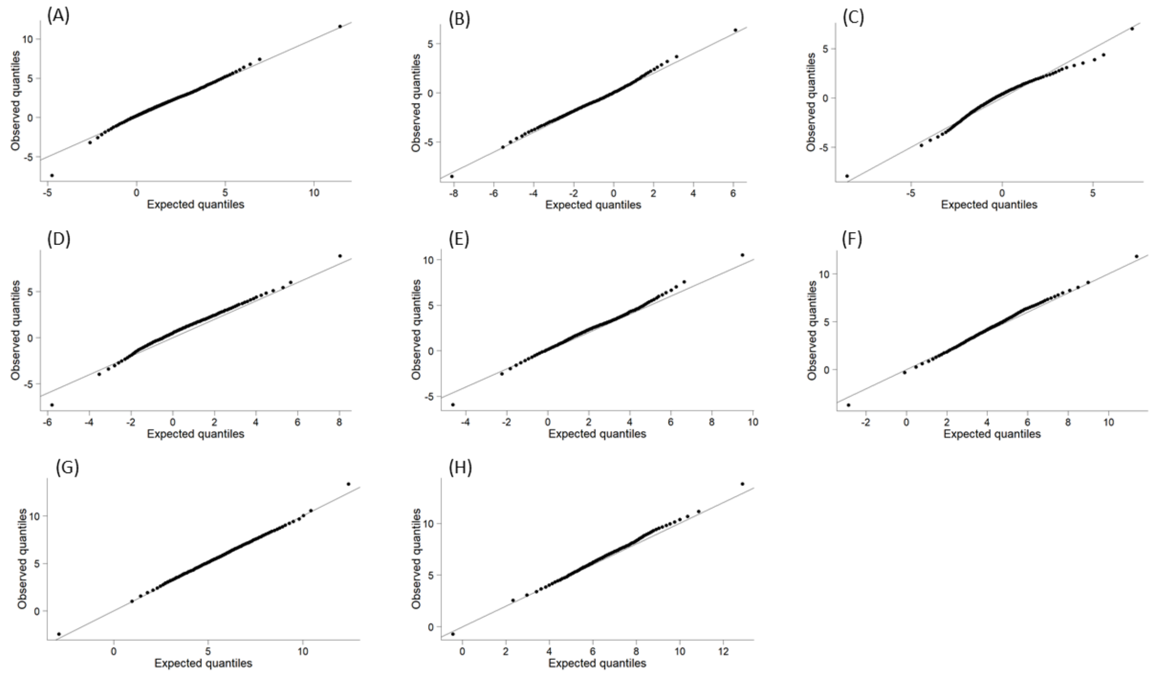

**Figure S5:** Q-Q plots for the model examining temporal and geographic trends in the minimum temperature during the observed incubation period for the Great Tit (A), and for the models examining temporal and geographic trends in the minimum temperature during fixed (null) windows for the Great Tit (B – H).

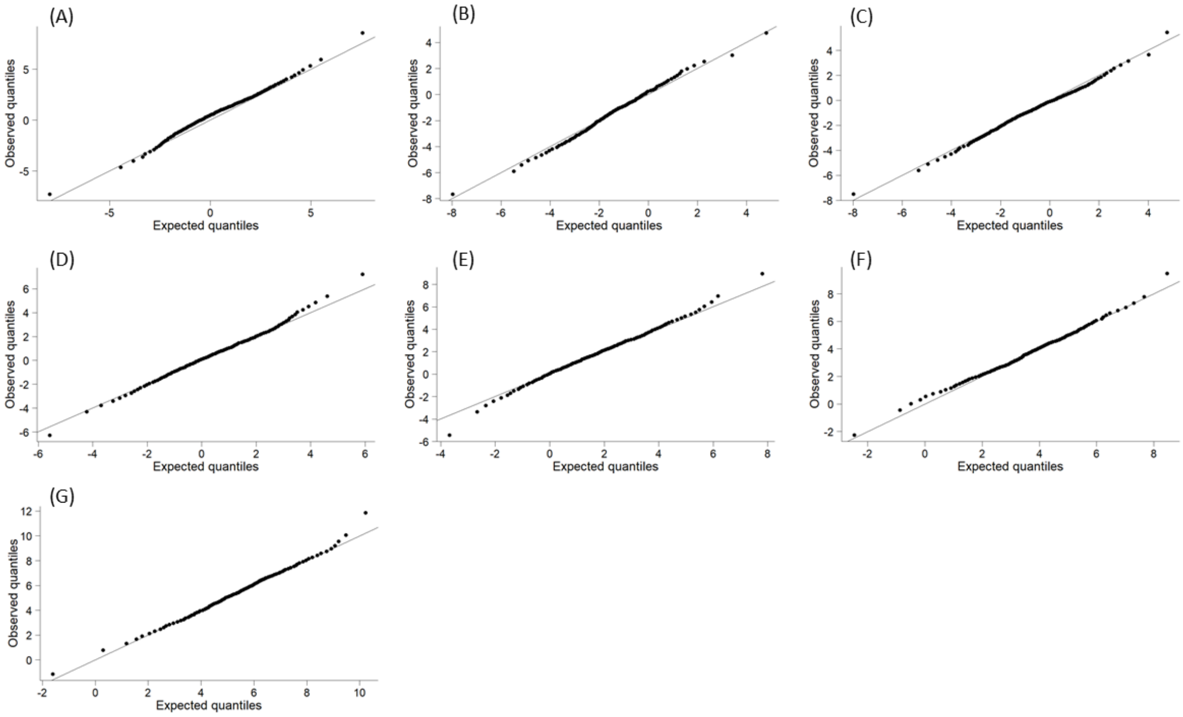

**Figure S6:** Q-Q plots for the model examining temporal and geographic trends in the minimum temperature during the observed incubation period for the Long-tailed Tit (A), and for the models examining temporal and geographic trends in the minimum temperature during fixed (null) windows for the Long-tailed Tit (B – G).

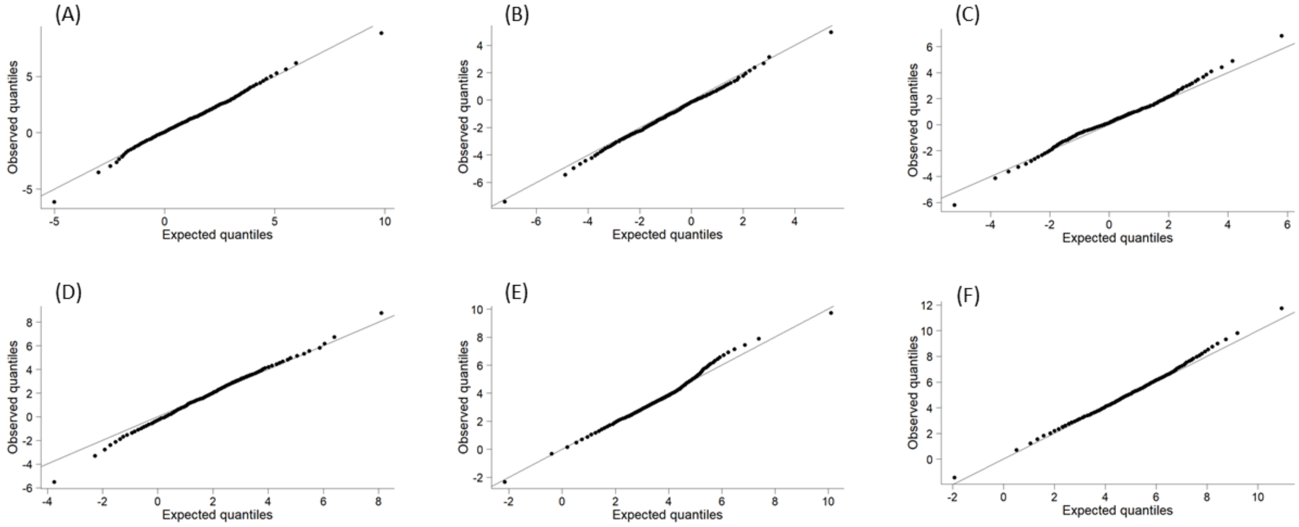

**Figure S7:** Q-Q plots for the model examining temporal and geographic trends in the minimum temperature during the observed incubation period for the Nuthatch (A), and for the models examining temporal and geographic trends in the minimum temperature during fixed (null) windows for the Nuthatch (B – F).

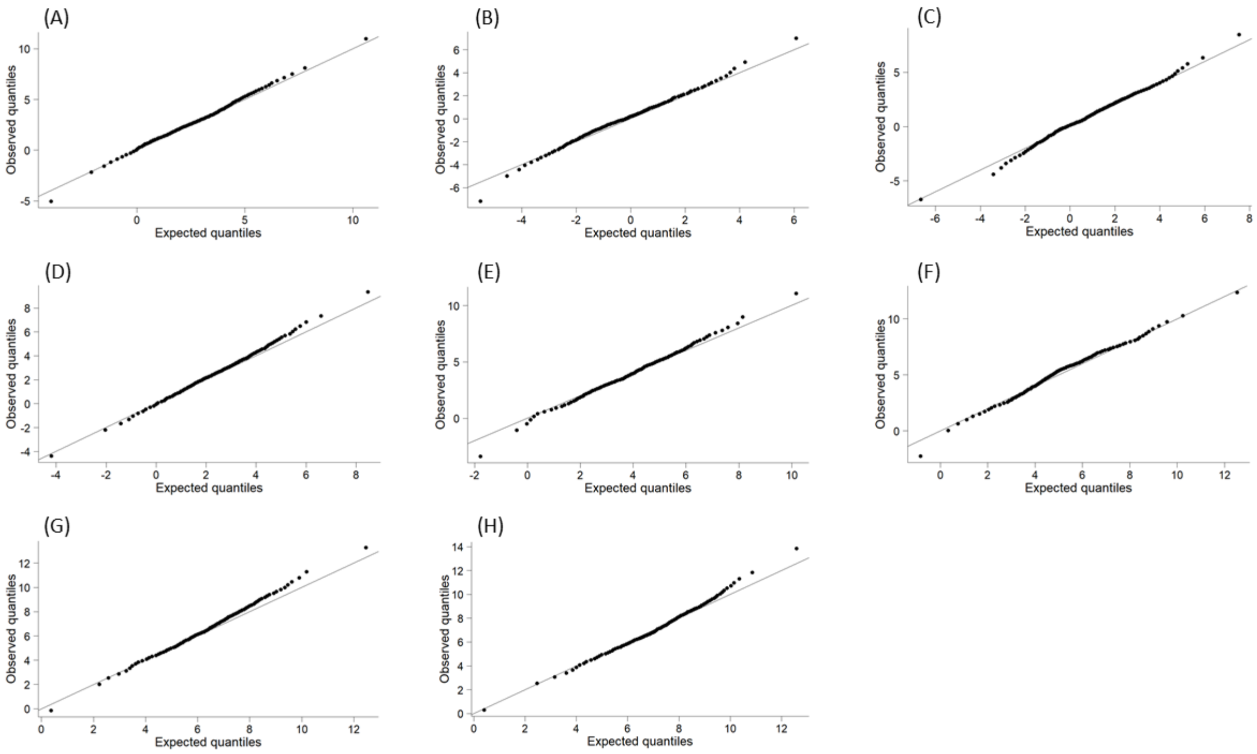

**Figure S8:** Q-Q plots for the model examining temporal and geographic trends in the minimum temperature during the observed incubation period for the Common Redstart (A), and for the models examining temporal and geographic trends in the minimum temperature during fixed (null) windows for the Common Redstart (B – H).

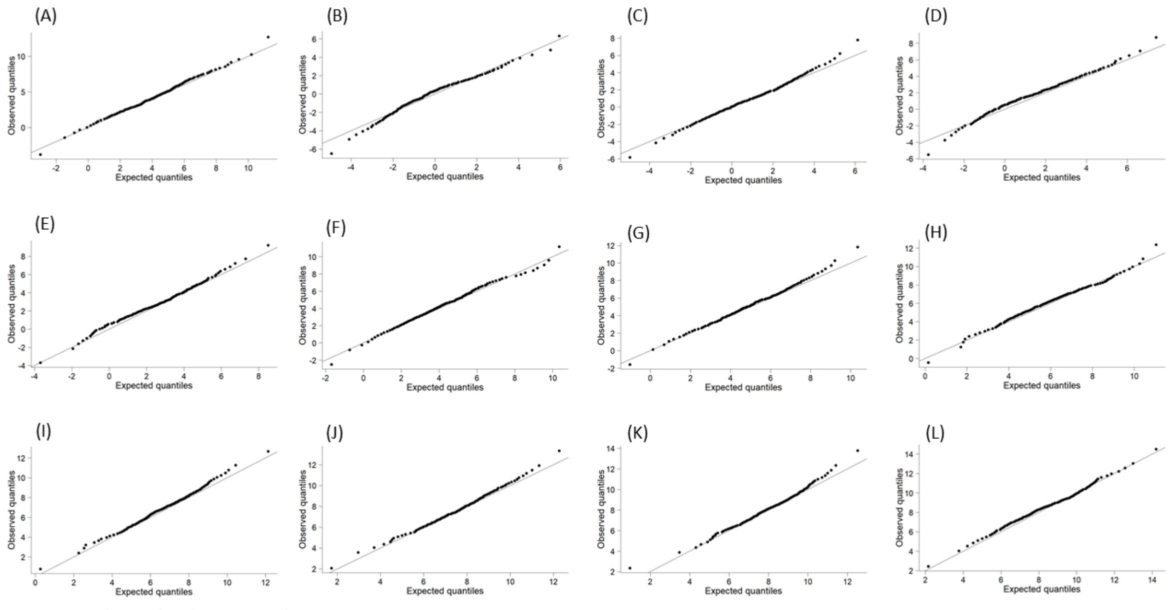

**Figure S9:** Q-Q plots for the model examining temporal and geographic trends in the minimum temperature during the observed incubation period for the Garden Warbler (A), and for the models examining temporal and geographic trends in the minimum temperature during fixed (null) windows for the Garden Warbler (B – L).

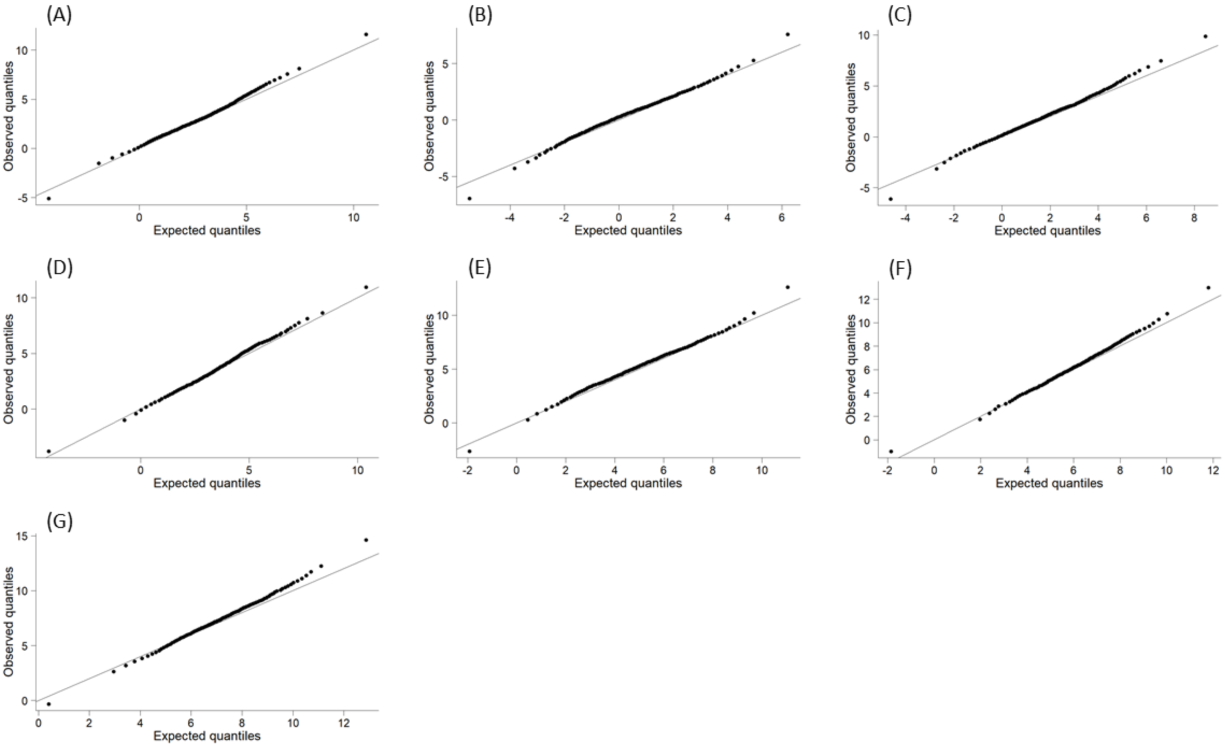

**Figure S10:** Q-Q plots for the model examining temporal and geographic trends in the minimum temperature during the observed incubation period for the Pied Flycatcher (A), and for the models examining temporal and geographic trends in the minimum temperature during fixed (null) windows for the Pied Flycatcher (B – G).

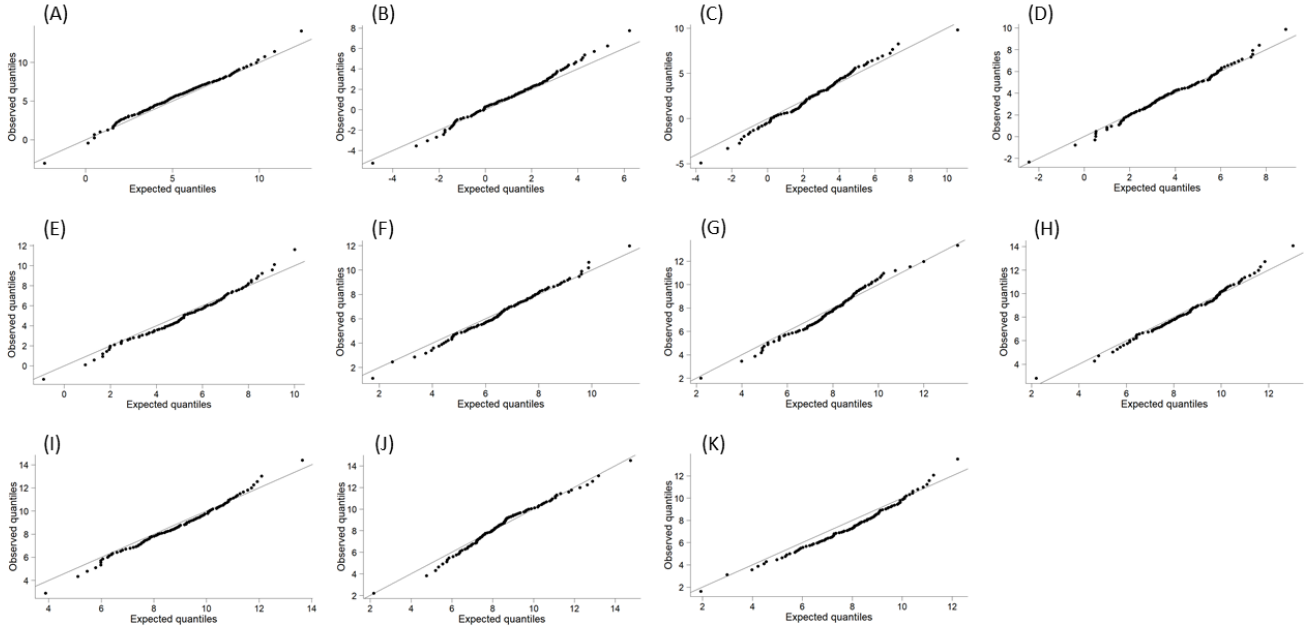

**Figure S11:** Q-Q plots for the model examining temporal and geographic trends in the minimum temperature during the observed incubation period for the Sedge Warbler (A), and for the models examining temporal and geographic trends in the minimum temperature during fixed (null) windows for the Sedge Warbler (B – K).

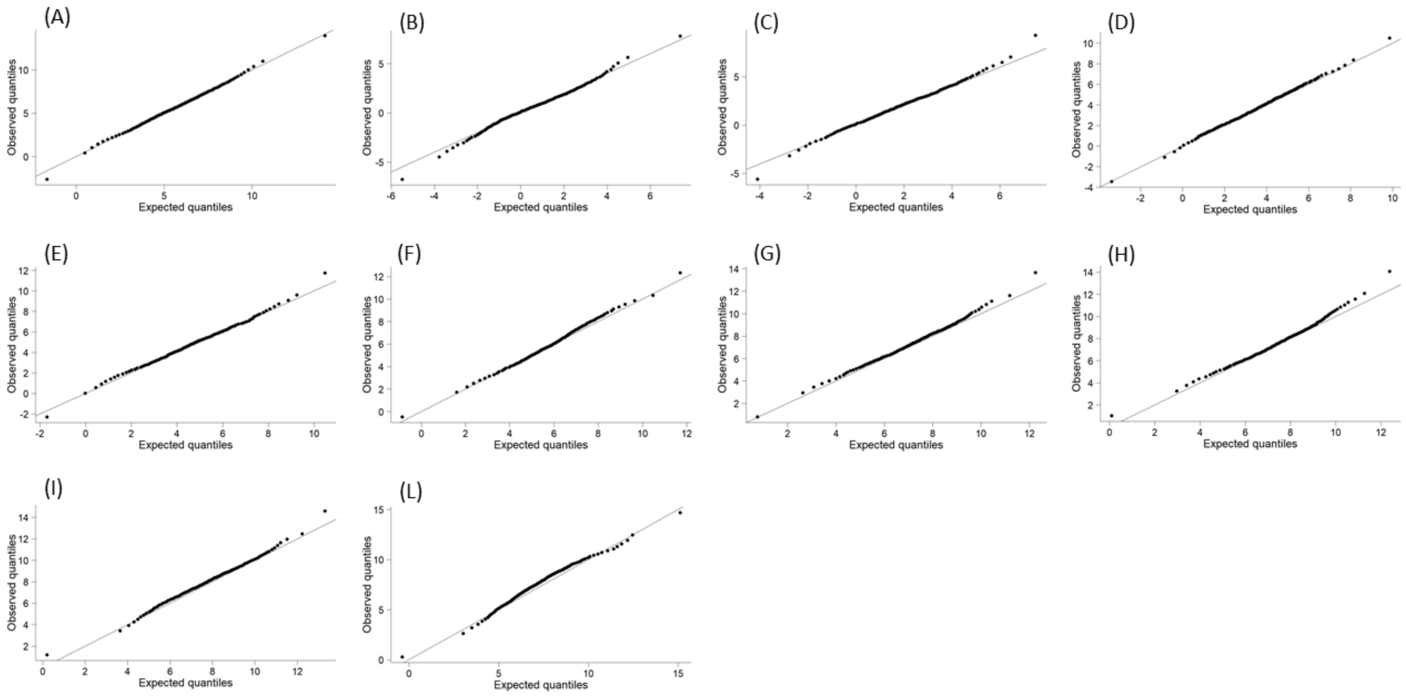

**Figure S12:** Q-Q plots for the model examining temporal and geographic trends in the minimum temperature during the observed incubation period for the Spotted Flycatcher (A), and for the models examining temporal and geographic trends in the minimum temperature during fixed (null) windows for the Spotted Flycatcher (B – L).

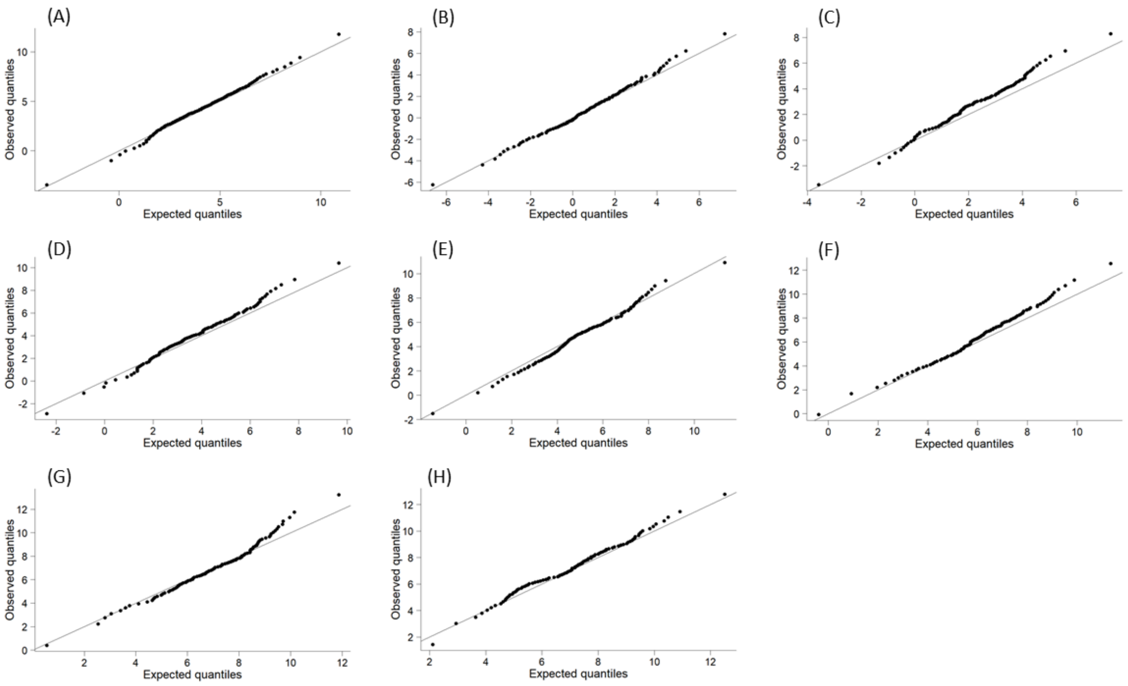

**Figure S13:** Q-Q plots for the model examining temporal and geographic trends in the minimum temperature during the observed incubation period for the Whinchat (A), and for the models examining temporal and geographic trends in the minimum temperature during fixed (null) windows for the Whinchat (B – H).

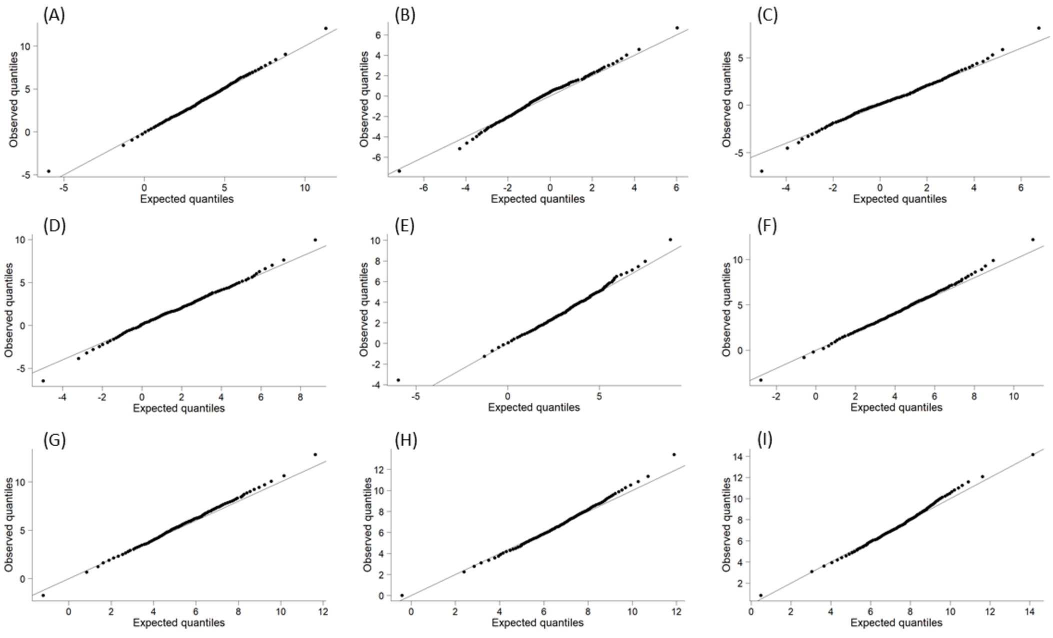

**Figure S14:** Q-Q plots for the model examining temporal and geographic trends in the minimum temperature during the observed incubation period for the Willow Warbler (A), and for the models examining temporal and geographic trends in the minimum temperature during fixed (null) windows for the Willow Warbler (B – I).

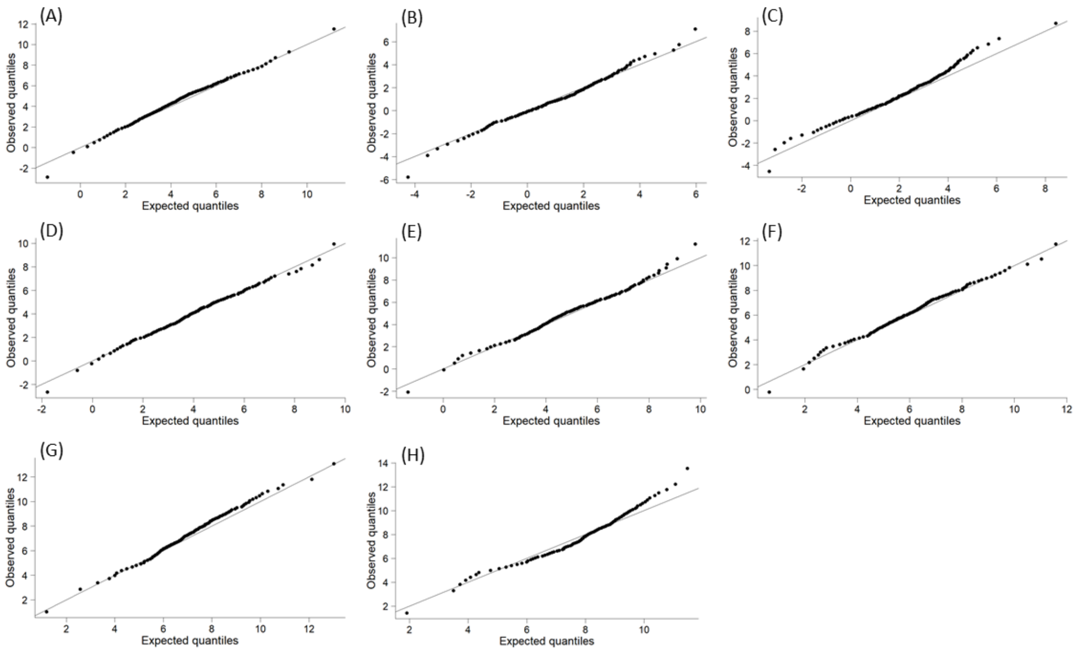

**Figure S15:** Q-Q plots for the model examining temporal and geographic trends in the minimum temperature during the observed incubation period for the Wood Warbler (A), and for the models examining temporal and geographic trends in the minimum temperature during fixed (null) windows for the Wood Warbler (B – H).

### Results

**Table S1:** Temporal years trends in thermal changes during incubation (Incubation trend) and in the null expectation of thermal changes (Average fixed window trend), along with the difference between these trends (Difference), are presented for five resident passerine species and eight migratory species across the UK over the past 40–50 years, based on BTO Nest Record Scheme data. Trends in the null expectation were assessed using minimum temperatures during fixed intervals (see Statistical Analysis in the main text for details). Average fixed window trends were calculated across the posterior estimates for these intervals. The Difference between the posterior slopes of the Incubation trend and the Average fixed window trend was used to test the null hypothesis of perfect thermal niche tracking. Coefficients with 95% credible intervals not overlapping 0 are highlighted in bold.

| | | **Incubation trend** | **Average fixed window trend** | **Difference (Incubation – Average fixed window)** |
|---|---|---|---|---|
| **Species** | **Species type** | **Year coefficient (l-95% CI, u-95% CI)** | **Year coefficient (l-95% CI, u-95% CI)** | **Year coefficient (l-95% CI, u-95% CI)** |
| Blue Tit | Resident | -0.01 (-0.03, 0.01) | 0.03 (-0.01, 0.06) | 0.03 (-0.01, 0.07) |
| Chaffinch | | -0.001 (-0.02, 0.02) | 0.02 (-0.01, 0.06) | 0.03 (-0.01, 0.07) |
| Great Tit | | -0.01 (-0.03, 0.01) | 0.03 (-0.01, 0.06) | 0.04 (-0.0002, 0.08) |
| Long-tailed Tit | | -0.01 (-0.03, 0.02) | 0.03 (-0.01, 0.06) | 0.04 (-0.01, 0.08) |
| Nuthatch | | -0.004 (-0.03, 0.02) | 0.02 (-0.01, 0.05) | 0.03 (-0.01, 0.07) |
| Garden Warbler | Migratory | 0.01 (-0.01, 0.04) | 0.03 (-0.01, 0.07) | 0.01 (-0.03, 0.06) |
| Pied Flycatcher | | 0.005 (-0.02, 0.03) | 0.02 (-0.01, 0.06) | 0.02 (-0.02, 0.06) |
| Common Redstart | | 0.005 (-0.02, 0.03) | 0.02 (-0.03, 0.07) | 0.02 (-0.04, 0.07) |
| Sedge Warbler | | 0.02 (-0.01, 0.04) | 0.03 (-0.02, 0.07) | 0.01 (-0.04, 0.06) |
| Spotted Flycatcher | | **0.03 (0.01, 0.04)** | 0.02 (-0.02, 0.06) | -0.001 (-0.05, 0.04) |
| Whinchat | | **0.02 (0.001, 0.04)** | 0.03 (-0.01, 0.06) | 0.004 (-0.03, 0.04) |
| Willow Warbler | | 0.01 (-0.01, 0.03) | 0.02 (-0.01, 0.05) | 0.01 (-0.03, 0.05) |
| Wood Warbler | | 0.02 (-0.004, 0.04) | 0.02 (-0.02, 0.06) | 0.007 (-0.04, 0.05) |

**Table S2:** Interannual trends in thermal changes during incubation (Incubation trend) and in the null expectation of thermal changes (Average fixed window trend), along with the difference between these trends (Difference), are presented for five resident passerine species and eight migratory species across the UK over the past 40–50 years, based on BTO Nest Record Scheme data. Trends in the null expectation were assessed using minimum temperatures during fixed intervals (see Statistical Analysis in the main text for details). Average fixed window trends were calculated across the posterior estimates for these intervals. The Difference between the posterior slopes of the Incubation trend and the Average fixed window trend was used to test the null hypothesis of perfect thermal niche tracking. Coefficients with 95% credible intervals not overlapping 0 are highlighted in bold.

| | | Incubation trend | Average fixed window trend | Difference (Incubation – Average fixed window) |
|---|---|---|---|---|
| **Species** | **Species type** | **Interannual coefficient (l-95% CI, u-95% CI)** | **Interannual coefficient (l-95% CI, u-95% CI)** | **Interannual coefficient (l-95% CI, u-95% CI)** |
| Blue Tit | Resident | **1.28 (0.83, 1.91)** | **2.32 (1.04, 4.03)** | 1.05 (-0.37, 2.82) |
| Chaffinch | | **0.72 (0.46, 1.09)** | **2.2 (0.95, 3.83)** | **1.5 (0.16, 3.14)** |
| Great Tit | | **1.06 (0.7, 1.58)** | **2.07 (0.94, 3.39)** | 1.01 (-0.21, 2.39) |
| Long-Tailed Tit | | **1.23 (0.78, 1.89)** | **2.17 (0.84, 3.85)** | 0.95 (-0.58, 2.72) |
| Nuthatch | | **1.28 (0.83, 1.96)** | **2.08 (0.98, 3.83)** | 0.8 (-0.51, 2.64) |
| Garden Warbler | Migratory | **0.76 (0.43, 1.29)** | **2.72 (1.21, 4.87)** | **1.95 (0.33, 4.12)** |
| Pied Flycatcher | | **1.35 (0.88, 2.04)** | **2.17 (1.19, 3.68)** | 0.83 (-0.38, 2.42) |
| Common Redstart | | **0.99 (0.62, 1.54)** | **2.24 (1.22, 4.12)** | **1.25 (0.07, 3.16)** |
| Sedge Warbler | | **0.71 (0.38, 1.23)** | **2.06 (0.93, 4.88)** | **1.35 (0.07, 4.2)** |
| Spotted Flycatcher | | **0.52 (0.32, 0.82)** | **1.91 (0.91, 3.53)** | **1.39 (0.33, 3.0)** |
| Whinchat | | **0.72 (0.4, 1.2)** | **1.98 (0.88, 3.93)** | **1.26 (0.05, 3.24)** |
| Willow Warbler | | **0.91 (0.58, 1.42)** | **2.28 (1.05, 4.46)** | **1.37 (0.01, 3.57)** |
| Wood Warbler | | **0.9 (0.54, 1.45)** | **2.09 (1.14, 3.52)** | **1.18 (0.08, 2.68)** |

**Table S3:** Spatial latitudinal trends in thermal changes during incubation (Incubation trend) and in the null expectation of thermal changes (Average fixed window trend), along with the difference between these trends (Difference), are presented for five resident passerine species and eight migratory species across the UK over the past 40–50 years, based on BTO Nest Record Scheme data. Trends in the null expectation were assessed using minimum temperatures during fixed intervals (see Statistical Analysis in the main text for details). Average fixed window trends were calculated across the posterior estimates for these intervals. The Difference between the posterior slopes of the Incubation trend and the Average fixed window trend was used to test the null hypothesis of perfect thermal niche tracking. Coefficients with 95% credible intervals not overlapping 0 are highlighted in bold.

| | | **Incubation trend** | **Average fixed window trend** | **Difference (Incubation – Average fixed window)** |
|---|---|---|---|---|
| **Species** | **Species type** | **Latitude coefficient (l-95% CI, u-95% CI)** | **Latitude coefficient (l-95% CI, u-95% CI)** | **Latitude coefficient (l-95% CI, u-95% CI)** |
| Blue Tit | Resident | **-0.18 (-0.24, -0.12)** | **-0.33 (-0.41, -0.2)** | **-0.15 (-0.25, -0.001)** |
| Chaffinch | | **-0.17 (-0.24, -0.11)** | **-0.31 (-0.43, -0.16)** | -0.14 (-0.28, 0.03) |
| Great Tit | | **-0.13 (-0.2, -0.07)** | **-0.32 (-0.41, -0.2)** | **-0.19 (-0.31, -0.05)** |
| Long-tailed Tit | | **-0.2 (-0.32, -0.08)** | **-0.25 (-0.43, -0.1)** | -0.06 (-0.27, 0.13) |
| Nuthatch | | -0.1 (-0.22, 0.02) | **-0.34 (-0.46, -0.23)** | **-0.24 (-0.41, -0.08)** |
| Garden Warbler | Migratory | **-0.23 (-0.4, -0.06)** | **-0.27 (-0.42, -0.12)** | -0.03 (-0.26, 0.2) |
| Pied Flycatcher | | **-0.26 (-0.35, -0.16)** | **-0.46 (-0.59, -0.35)** | **-0.2 (-0.36, -0.05)** |
| Common Redstart | | **-0.29 (-0.37, -0.21)** | **-0.36 (-0.46, -0.26)** | -0.07 (-0.2, 0.06) |
| Sedge Warbler | | 0.003 (-0.18, 0.10) | **-0.24 (-0.37, -0.11)** | -0.24 (-0.43, 0.06) |
| Spotted Flycatcher | | **-0.33 (-0.4, -0.27)** | **-0.34 (-0.43, -0.24)** | -0.01 (-0.12, 0.11) |
| Whinchat | | **-0.4 (-0.51, -0.29)** | **-0.34 (-0.46, -0.22)** | 0.06 (-0.1, 0.22) |
| Willow Warbler | | **-0.19 (-0.26, -0.12)** | **-0.29 (-0.38, -0.16)** | -0.1 (-0.22, 0.05) |
| Wood Warbler | | **-0.3 (-0.41, -0.18)** | **-0.38 (-0.51, -0.22)** | -0.09 (-0.26, 0.11) |

**Table S4:** Spatial elevational trends in thermal changes during incubation (Incubation trend) and in the null expectation of thermal changes (Average fixed window trend), along with the difference between these trends (Difference), are presented for five resident passerine species and eight migratory species across the UK over the past 40–50 years, based on BTO Nest Record Scheme data. Trends in the null expectation were assessed using minimum temperatures during fixed intervals (see Statistical Analysis in the main text for details). Average fixed window trends were calculated across the posterior estimates for these intervals. The Difference between the posterior slopes of the Incubation trend and the Average fixed window trend was used to test the null hypothesis of perfect thermal niche tracking. Coefficients with 95% credible intervals not overlapping 0 are highlighted in bold.

| | | **Incubation trend** | **Average fixed window trend** | **Difference (Incubation – Average fixed window)** |
|---|---|---|---|---|
| **Species** | **Species type** | **Elevation coefficient (l-95% CI, u-95% CI)** | **Elevation coefficient (l-95% CI, u-95% CI)** | **Elevation coefficient (l-95% CI, u-95% CI)** |
| Blue Tit | Resident | **-0.002 (-0.002, -0.002)** | **-0.003, (-0.004, -0.003)** | **-0.001 (-0.002, -0.001)** |
| Chaffinch | | **-0.002 (-0.003, -0.0005)** | **-0.003 (-0.004, -0.002)** | -0.001 (-0.003, 0.0001) |
| Great Tit | | **-0.002 (-0.002, -0.001)** | **-0.003, (-0.004, -0.003)** | **-0.001 (-0.002, -0.001)** |
| Long-tailed Tit | | -0.002 (-0.003, 0.00004) | **-0.004 (-0.005, -0.002)** | -0.002 (-0.004, 0.0002) |
| Nuthatch | | **-0.002 (-0.003, -0.001)** | **-0.003 (-0.004, -0.003)** | **-0.001 (-0.003, -0.0002)** |
| Garden Warbler | Migratory | **-0.004 (-0.01, -0.002)** | **-0.004 (-0.005, -0.003)** | 0.0003 (-0.002, 0.003) |
| Pied Flycatcher | | **-0.002 (-0.003, -0.002)** | **-0.003 (-0.004, -0.003)** | **-0.001 (-0.002, -0.0003)** |
| Common Redstart | | **-0.003 (-0.004, -0.002)** | **-0.004 (-0.004, -0.003)** | -0.0004 (-0.001, 0.001) |
| Sedge Warbler | | **-0.004 (-0.01, -0.0004)** | **-0.002 (-0.005, 0.0002)** | 0.001 (-0.002, 0.005) |
| Spotted Flycatcher | | **-0.005 (-0.006, -0.004)** | **-0.004 (-0.005, -0.003)** | **0.002 (0.0004, 0.003)** |
| Whinchat | | **-0.004 (-0.005, -0.003)** | **-0.004 (-0.005, -0.004)** | -0.00004 (-0.001, 0.001) |
| Willow Warbler | | **-0.004 (-0.005, -0.003)** | **-0.004 (-0.005, -0.004)** | -0.0005 (-0.001, 0.001) |
| Wood Warbler | | **-0.002 (-0.003, -0.0002)** | **-0.003 (-0.004, -0.002)** | -0.001 (-0.003, 0.0003) |

**Table S5:** Thermal niche tracking metrics across years, interannual variations (year random effect), latitudes and elevations on thirteen passerine species (names reported as BTO codes) across the UK for the past 40-50 years. The metric is calculated as the ratio between observed trends in incubation temperatures and trends in fixed-window temperatures. A value of 0 indicates perfect thermal niche tracking, while a value of 1 indicates no tracking. Reported coefficients are the median tracking metric across the posterior for each temporal and spatial dimension. Coefficients and credible intervals (95% CIs) that do not overlap 0 or 1 are highlighted in bold.

| Species | Species type | Year coefficient (l-95% CI, u-95% CI) | Interannual coefficient (l-95% CI, u-95% CI) | Latitude coefficient (l-95% CI, u-95% CI) | Elevation coefficient (l-95% CI, u-95% CI) |
|---|---|---|---|---|---|
| Blue Tit | Resident | 0.41 (0.02, 6.89) | 0.56 (0.27, 1.32) | **0.54 (0.35, 0.99)** | **0.59 (0.47, 0.73)** |
| Chaffinch | | 0.26 (0.01, 5.21) | **0.32 (0.16, 0.84)** | 0.55 (0.30, 1.17) | 0.54 (0.17, 1.02) |
| Great Tit | | 0.47 (0.02, 8.1) | 0.51 (0.27, 1.21) | **0.40 (0.20, 0.76)** | **0.57 (0.42, 0.75)** |
| Long-Tailed Tit | | 0.37 (0.02, 5.76) | 0.56 (0.26, 1.61) | 0.79 (0.27, 2.04) | 0.45 (0.05, 1.06) |
| Nuthatch | | 0.36 (0.02, 6.01) | 0.63 (0.29, 1.46) | **0.30 (0.02, 0.71)** | **0.57 (0.26, 0.93)** |
| Garden Warbler | Migratory | 0.49 (0.02, 9.31) | **0.29 (0.12, 0.75)** | 0.87 (0.22, 2.34) | 1.10 (0.43, 1.94) |
| Pied Flycatcher | | 0.37 (0.01, 6.68) | 0.63 (0.31, 1.28) | **0.56 (0.33, 0.87)** | **0.70 (0.54, 0.88)** |
| Common Redstart | | 0.40 (0.02, 9.39) | **0.46 (0.20, 0.95)** | 0.81 (0.55, 1.21) | **0.46 (0.20, 0.95)** |
| Sedge Warbler | | 0.54 (0.03, 9.87) | **0.38 (0.12, 0.93)** | **0.19 (0.01, 0.78)** | **0.38 (012, 0.93)** |
| Spotted Flycatcher | | 0.99 (0.24, 17.09) | **0.28 (0.12, 0.67)** | 0.94 (0.70, 1.42) | 1.50 (1.10, 2.05) |
| Whinchat | | 0.84 (0.11, 12.08) | **0.38 (0.15, 0.95)** | 1.18 (0.77, 1.91) | 0.99 (0.67, 1.33) |
| Willow Warbler | | 0.51 (0.02, 10.36) | **0.42 (0.18, 0.99)** | 0.65 (0.38, 1.30) | **0.42 (0.18, 0.99)** |
| Wood Warbler | | 0.69 (0.04, 15.02) | 0.44 (0.20, 0.94) | 0.77 (0.44, 1.44) | **0.44 (0.20, 0.94)** |

**Table S6:** Pairwise comparisons between thermal niche tracking metrics across years, interannual variations, latitudes and elevations on 13 passerines species (All species: five resident species and eight migratory species) and on 9 passerine species (Exclusion of second brood species: five resident species and four migratory species) across the UK for the past 40-50 years. The species excluded for showing signs of potential second broods were Common Redstart, Garden Warbler, Spotted Flycatcher, and Willow Warbler. Reported coefficients represent the median difference of the tracking metric between the posterior distributions of the compared terms. Coefficients and 95% credible intervals (CIs) that do not overlap 0 or 1 are highlighted in bold.

| | **All species** | | | **Exclusion of second brooded species** | | |
|---|---|---|---|---|---|---|
| **Average Posteriors Difference (A-B)** | **Median difference** | **l-95% CI difference** | **u-95% CI difference** | **Median difference** | **l-95% CI difference** | **u-95% CI difference** |
| Year — Latitude | -0.14 | -0.47 | 0.38 | -0.09 | -0.44 | 0.53 |
| Year — Elevation | -0.20 | -0.50 | 0.32 | -0.15 | -0.47 | 0.47 |
| Year — Interannual | 0.05 | -0.26 | 0.56 | -0.03 | -0.38 | 0.59 |
| Latitude — Elevation | -0.05 | -0.26 | 0.17 | -0.06 | -0.26 | 0.17 |
| Latitude — Interannual | 0.20 | -0.02 | 0.42 | 0.06 | -0.20 | 0.31 |
| Elevation — Interannual | **0.25** | **0.05** | **0.44** | 0.12 | -0.11 | 0.32 |

**Table S7:** Latitude and elevation ranges for thirteen passerine species in the UK. Minimum and maximum values are reported, with the interquartile range shown in brackets.

| **Species** | **Latitude (°North) min – max ($25^{th}$ percentile – $75^{th}$ percentile)** | **Elevation (metres) min – max ($25^{th}$ percentile – $75^{th}$ percentile)** |
|---|---|---|
| Blue tit | 50.09 – 58.48 (51.57 – 53.30) | 0 – 604 (49 – 156) |
| Chaffinch | 50.10 – 58.98 (51.70 – 53.65) | 0 – 635 (38 – 127) |
| Great Tit | 50.09 – 58.53 (51.67 – 53.86) | 0 – 736 (47 – 149) |
| Long-tailed Tit | 50.10 – 57.17 (51.62 – 53.53) | 0 – 369 (29 – 106) |
| Nuthatch | 50.11 – 56.25 (51.12 – 53.14) | 4 – 550 (86 – 198) |
| Common Redstart | 50.42 – 58.41 (51.88 – 54.52) | 1 – 667 (135 – 255) |
| Garden Warbler | 50.17 – 57.34 (51.78 – 53.23) | 1 – 463 (61 – 140) |
| Pied Flycatcher | 50.24 – 57.90 (51.86 – 54.08) | 1 – 771 (135 – 232) |
| Sedge Warbler | 50.13 – 60.05 (51.77 – 53.39) | 0 – 461 (17 – 41.5) |
| Spotted Flycatcher | 50.22 – 58.49 (51.83 – 54.34) | 0 – 514 (45 – 160) |
| Whinchat | 50.43 – 59.07 (51.24 – 54.90) | 1 – 609 (264 – 382) |
| Willow Warbler | 50.17 – 60.15 (51.35 – 54.54) | 0 – 812 (64 – 196) |
| Wood Warbler | 50.48 – 58.48 (50.91– 52.76) | 0 – 566 (126 – 250) |

**Table S8:** Hatching probability trends in response to minimum incubation temperature (both linear and quadratic effect), laying date deviations (both linear and quadratic effect), and year are presented for thirteen passerine species across the UK over the past 40–50 years, based on BTO Nest Record Scheme data. Hatching probability was treated as a binary response, with nests containing at least one live chick at least on day 5 days post-hatching classified as successful (nest success = 1) and nests with no alive chicks at that stage considered unsuccessful (nest success = 0). Laying date deviations were calculated as the difference between the observed laying date for a nest and the annual mean laying date within the corresponding 1 km grid cell. The fixed effect of year was scaled for the analyses on Blue tits (Schielzeth, 2010), but we here report the re-scaled slope and 95% credible intervals (CIs). Coefficients are expressed in logit units. Coefficients with 95% CIs not overlapping 0 are highlighted in bold.

| **Spotted Flycatcher** | **Whinchat** | **Willow Warbler** | **Wood Warbler** |
|---|---|---|---|
| coefficient (l-95% CI, u-95% CI) | coefficient (l-95% CI, u-95% CI) | coefficient (l-95% CI, u-95% CI) | coefficient (l-95% CI, u-95% CI) |
| 7.65 (-22.3, 33.01) | 41.34 (-5.39, 85.26) | **46.95 (19.46, 72.0)** | 26.9 (-11.5, 67.05) |
| 0.05 (-0.11, 0.24) | 0.001 (-0.24, 0.24) | 0.09 (-0.01, 0.19) | 0.006 (-0.22, 0.22) |
| 0.0005 (-0.01, 0.02) | -0.01 (-0.04, 0.01) | -0.01 (-0.03, 0.0003) | -0.01 (-0.03, 0.01) |
| 0.0004 (-0.02, 0.02) | **-0.07 (-0.1, -0.008)** | **-0.05 (-0.08, -0.02)** | -0.03 (-0.08, 0.02) |
| **0.002 (0.001, 0.002)** | -0.00005 (-0.001, 0.001) | **0.003 (0.002, 0.005)** | 0.0001 (-0.001, 0.002) |
| -0.003 (-0.02, 0.01) | -0.02 (-0.04, 0.004) | **-0.02 (-0.03, -0.009)** | -0.01 (-0.03, 0.006) |
| **variance (l-95% CI, u-95% CI)** | **variance (l-95% CI, u-95% CI)** | **variance (l-95% CI, u-95% CI)** | **variance (l-95% CI, u-95% CI)** |
| 0.12 (0.05, 0.21) | 0.37 (0.1, 0.73) | 0.21 (0.09, 0.34) | 0.2 (0.06, 0.37) |
| 0.12 (0.05, 0.21) | 0.43 (0.09, 0.86) | 0.23 (0.08, 0.4) | 0.40 (0.08, 0.78) |
| 0.99 (0.57, 1.45) | 1.05 (0.43, 1.75) | 0.45 (0.24, 0.68) | 0.74 (0.32, 1.21) |
| 0.003 (-0.02, 0.03) | 0.03 (-0.06, 0.12) | 0.0007 (-0.03, 0.03) | 0.007 (-0.05, 0.06) |
| 0.02 (0.02, 0.02) | 0.06 (0.04, 0.08) | 0.04 (0.03, 0.05) | 0.05 (0.03, 0.07) |
| 0.78 (0.43, 1.15) | 0.48 (0.12, 0.96) | 0.64 (0.36, 0.98) | 0.63 (0.21, 1.12) |

| Great Tit | Long-Tailed Tit | Nuthatch | Garden Warbler | Pied Flycatcher | Redstart | Sedge Warbler |
|---|---|---|---|---|---|---|
| coefficient (l-95% CI, u-95% CI) | coefficient (l-95% CI, u-95% CI) | coefficient (l-95% CI, u-95% CI) | coefficient (l-95% CI, u-95% CI) | coefficient (l-95% CI, u-95% CI) | coefficient (l-95% CI, u-95% CI) | coefficient (l-95% CI, u-95% CI) |
| -4.14 (-0.25, 0.15) | -13.95 (-41.27, 14.88) | -1.96 (-43.56, 38.61) | 58.16 (20.04, 10.11) | **25.73 (2.84, 51.1)** | 2.81 (-28.2, 33.53) | 38.34 (-8.83, 85.95) |
| **-0.05 (-0.84, -0.01)** | **-0.07 (-0.14, -0.01)** | -0.02 (-0.14, 0.1) | -0.07 (-0.27, 0.11) | -0.05 (-0.1, 0.01) | -0.11 (-0.25, 0.03) | -0.11 (-0.36, 0.14) |
| 0.001 (-0.005, 0.008) | -0.005 (-0.02, 0.01) | -0.005 (-0.03, 0.03) | 0.008 (-0.01, 0.03) | -0.0003 (-0.01, 0.01) | 0.009 (-0.01, 0.03) | 0.01 (-0.01, 0.03) |
| **-0.06 (-0.07, -0.05)** | -0.01 (-0.06, 0.04) | **-0.08 (-0.14, -0.01)** | 0.006 (-0.07, 0.08) | **-0.06 (-0.07, -0.04)** | **-0.05 (-0.1, -0.02)** | **-0.06 (-0.09, -0.01)** |
| 0.0002 (-0.0004, 0.001) | **0.005 (0.001, 0.009)** | 0.001 (-0.005, 0.007) | **0.002 (0.0001, 0.004)** | **-0.002 (-0.003, -0.001)** | **0.005 (0.003, 0.008)** | **0.001 (0.0001, 0.002)** |
| 0.003 (-0.006, 0.01) | 0.007 (-0.007, 0.02) | 0.003 (-0.02, 0.02) | **-0.03 (-0.05, -0.009)** | -0.01 (-0.02, 0.0007) | 0.00006 (-0.01, 0.02) | -0.02 (-0.04, 0.005) |
| **variance (l-95% CI, u-95% CI)** | **variance (l-95% CI, u-95% CI)** | **variance (l-95% CI, u-95% CI)** | **variance (l-95% CI, u-95% CI)** | **variance (l-95% CI, u-95% CI)** | **variance (l-95% CI, u-95% CI)** | **variance (l-95% CI, u-95% CI)** |
| 0.10 (0.05, 0.15) | 0.14 (0.06, 0.24) | 0.32 (0.11, 0.59) | 0.25 (0.07, 0.49) | 0.16 (0.08, 0.26) | 0.13 (0.05, 0.22) | 0.322 (0.09, 0.6) |
| 0.08 (0.05, 0.13) | 0.32 (0.11, 0.55) | 0.18 (0.06, 0.34) | 0.5 (0.12, 0.97) | 0.19 (0.08, 0.34) | 0.20 (0.07, 0.39) | 0.58 (0.13, 1.18) |
| 0.29 (0.23, 0.36) | 0.59 (0.3, 0.96) | 0.73 (0.26, 1.21) | 0.57 (0.18, 1.07) | 0.73 (0.59, 0.88) | 0.71 (0.31, 1.12) | 0.59 (0.2, 1.06) |
| -0.01 (-0.01, -0.002) | 0.01 (-0.04, 0.06) | -0.01 (-0.09) | -0.03 (-0.12, 0.04) | -0.005 (-0.02, 0.01) | 0.02 (-0.03, 0.07) | -0.0005 (-0.04, 0.04) |
| 0.01 (0.01, 0.01) | 0.06 (0.04, 0.08) | 0.08 (0.05, 0.11) | 0.08 (0.05, 0.12) | 0.01 (0.01, 0.02) | 0.05 (0.03, 0.06) | 0.04 (0.03, 0.05) |
| 0.42 (0.33, 0.52) | 0.69 (0.33, 1.13) | 0.54 (0.17, 0.99) | 1.09 (0.33, 1.91) | 0.67 (0.48, 0.88) | 0.42 (0.16, 0.73) | 0.64 (0.14, 1.3) |

| | Blue Tit | Chaffinch |
|---|---|---|
| **Fixed term** | coefficient (l-95% CI, u-95% CI) | coefficient (l-95% CI, u-95% CI) |
| Intercept | **2.74 (2.59, 2.88)** | 1.53 (-0.17, 0.20) |
| Min. incubation temperature | **0.04 (0.01, 0.07)** | 0.03 (-0.03, 0.08) |
| Min. incubation temperature (quadratic term) | **-0.01 (-0.01, -0.003)** | 0.002 (-0.07, 0.01) |
| Laying date deviations | **-0.07 (-0.08, -0.06)** | -0.007 (-0.02, 0.01) |
| Laying date deviations (quadratic term) | **-0.001 (-0.001, -0.0002)** | **0.001 (0.0001, 0.001)** |
| Year | -0.14 (-0.69, 0.46) | -0.0005 (-0.001, 0.01) |
| **Random term** | **variance (l-95% CI, u-95% CI)** | **variance (l-95% CI, u-95% CI)** |
| Year | 0.08 (0.04, 0.13) | 0.09 (0.04, 0.14) |
| Km50 | 0.10 (0.05, 0.15) | 0.21 (0.10, 0.34) |
| (Intercept) : (Intercept).year : km50 | 0.35 (0.29, 0.41) | 0.46 (0.27, 0.68) |
| Laying date deviations : (Intercept).year : km50 | -0.005 (-0.01, 0.001) | -0.001 (-0.02, 0.01) |
| Laying date deviations : FEDdeviations.year : km50 | 0.01 (0.01, 0.01) | 0.02 (0.01, 0.02) |
| Km5 | 0.60 (0.51, 0.71) | 0.43 (0.25, 0.63) |

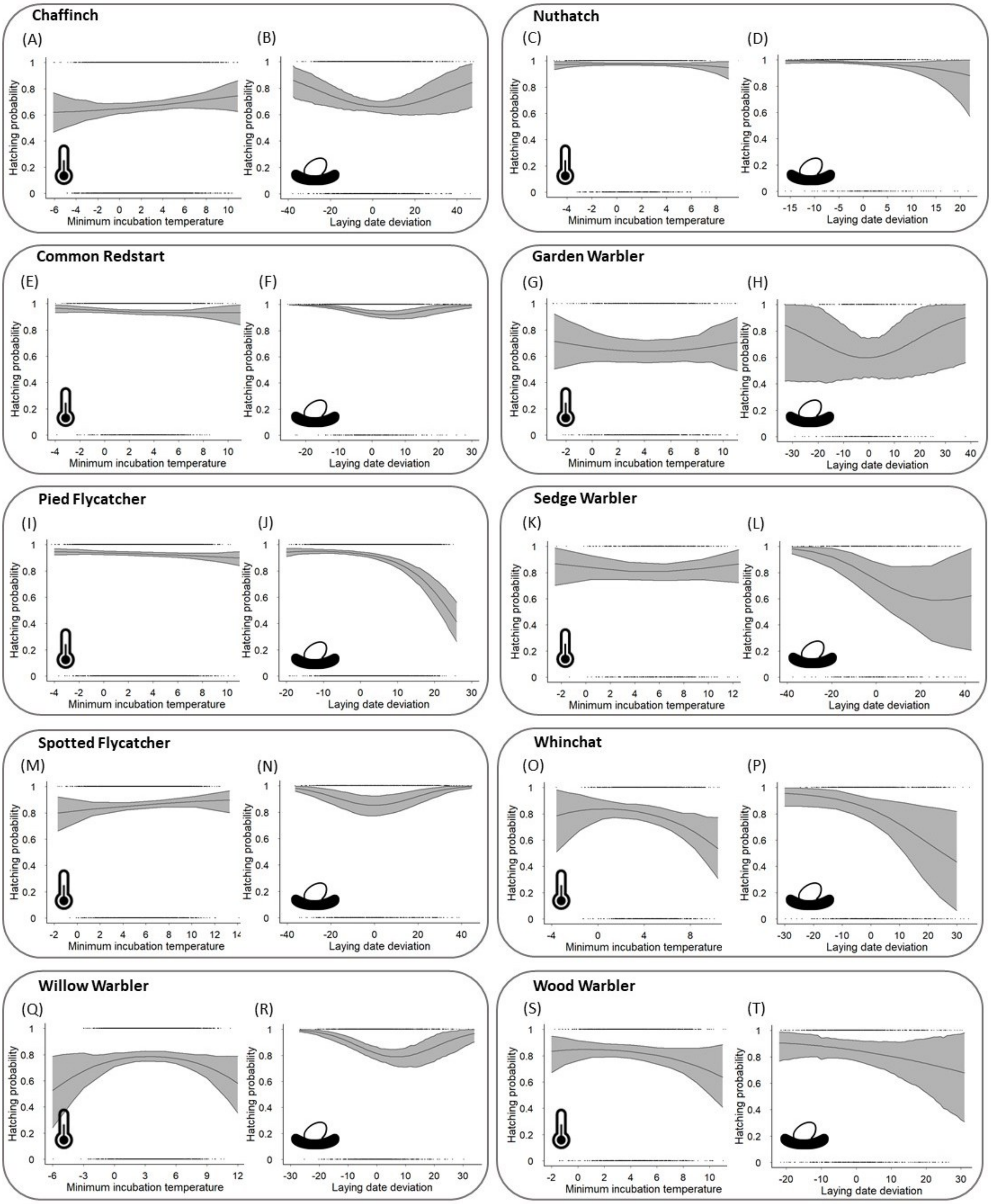

**Figure S16:** Trends in hatching probability in response to minimum incubation temperature and laying date deviations for ten passerine species, for the past 40-50 years, across the UK. Hatching probability was measured using a binary metric, with nests containing at least one alive chick at least on day 5 post-hatching classified as successful (hatching success = 1) and nests with no alive chicks considered unsuccessful (hatching success = 0). The trends slopes and 95% credible intervals are reported.

### References

Schielzeth, H. (2010) Simple means to improve the interpretability of regression coefficients. *Methods in Ecology and Evolution*, **1**, 103–113.
